# Combining Stability-Centered Atomistic Design with Machine Learning for Targeted Enzyme Optimization

**DOI:** 10.64898/2026.07.20.739108

**Authors:** Li Wan, Mahdi Bagherpoor Helabad, Lena Fraedrich, Sarel J. Fleishman, Martin J. Weissenborn

**Affiliations:** Institute of Chemistry, Martin Luther-University Halle-Wittenberg, Weinbergweg 22, 06120, Halle (Saale), Germany; Institute of Organic Chemistry, Leibniz University of Hannover, Schneiderberg 1B, 30167 Hannover, Germany; Department of Biomolecular Sciences, Weizmann Institute of Science, Rehovot 7600001, Israel

**Keywords:** Unspecific peroxygenase, Machine learning, Protein engineering, htFuncLib, β-damascone, oxyfunctionalization

## Abstract

FuncLib and high-throughput FuncLib (htFuncLib) generate diverse, functional protein libraries using a stability-centered design; however, this substrate-independent approach lacks target-specific functional constraints. We developed a machine-learning-assisted enzyme-engineering (MLEE) workflow that adds substrate-specific functional information to htFuncLib through an initial screening and sequencing round. The system was benchmarked using previously published four-position fitness landscapes of three different proteins. The MLEE workflow successfully generated compact libraries enriched in globally high-fitness variants. After the initial training phase, an MLEE-enriched library of just 12 variants increased the hit rate for the global top-0.05% variants by 5- to 12-fold relative to the htFuncLib baseline. Screening a larger set of 96 variants recovered at least one of these top-performing enzymes in 61.3–99.4% of the simulations. We then applied MLEE to *Mth*UPO-catalyzed β-damascone hydroxylation. Across two rounds, 506 distinct variants were screened and sequenced. While the initial substrate-independent htFuncLib library yielded 14% of variants with activity above the wild type, the MLEE-enriched library increased this hit rate to 90% (97 of 108 variants) with activity above the wild type. The best variant increased the turnover number for 4-hydroxy-β-damascone by 11.8-fold and achieved >99% regioisomeric excess. MLEE may bypass the need for transition-state models and reduce the effort required for obtaining high-activity variants.

**TABLE OF CONTENT:** 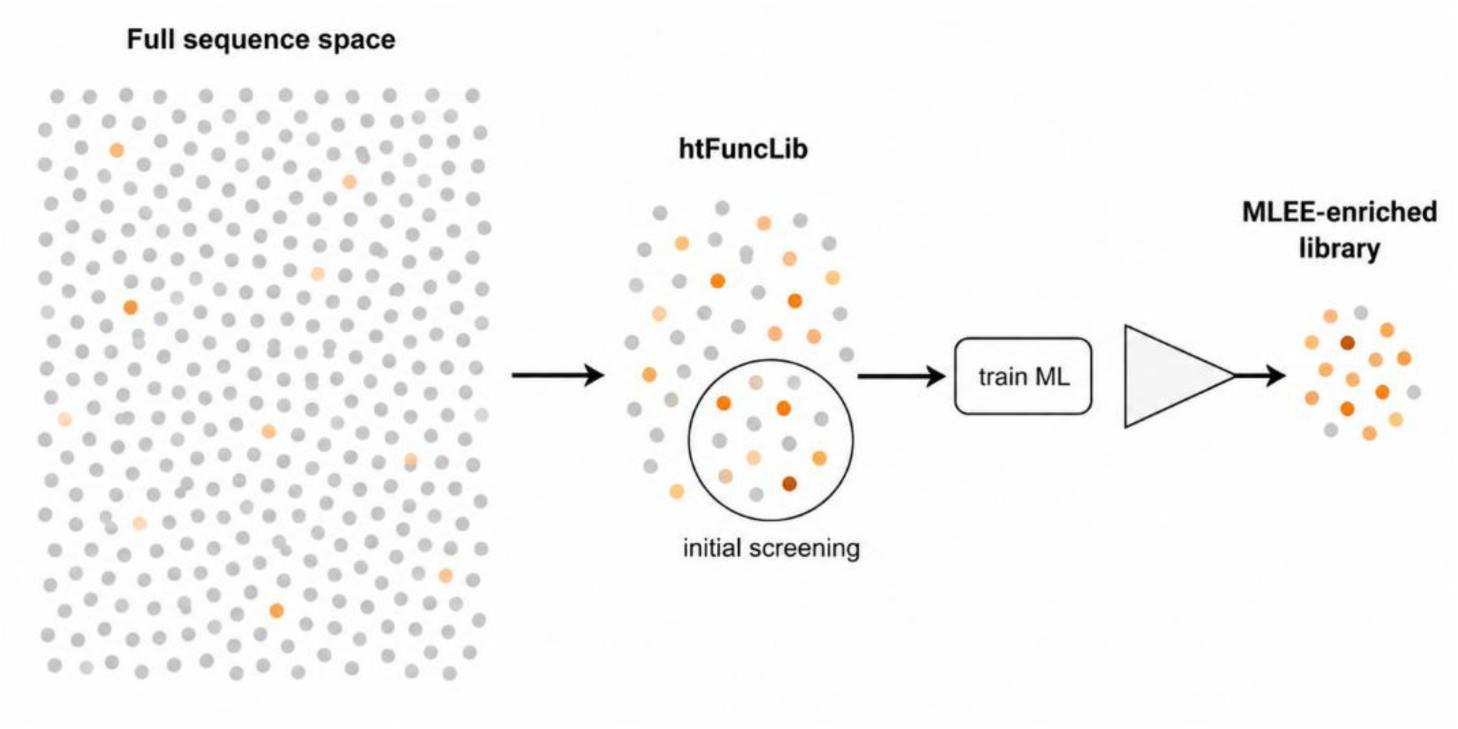

## Introduction

Engineering enzymes toward new or improved functions often requires coordinated mutation at several active-site positions. However, the number of possible combinations increases rapidly with each additional position, while many mutations are deleterious and their effects can change across different genetic backgrounds.^1–3^ Such epistatic interactions produce rugged fitness landscapes in which high-performing variants may be separated from the parent enzyme by low-fitness intermediates, limiting the efficiency of one-mutation-at-a-time optimization workflow used in directed evolution campaigns.^4–7^ Under experimentally limited screening capacity, which is a bottleneck faced by most enzyme engineering projects, the central challenge is therefore not only to reduce sequence space, but to define compact combinatorial libraries that retain mutational diversity while enriching for functional variants.^8–10^

FuncLib addresses this challenge by combining phylogenetic information with Rosetta native-state energy calculations to design multipoint variants predicted to be stable.^11^ This approach has generated diverse functional enzyme repertoires and enabled substantial improvements in several enzyme engineering projects.^12–14^ FuncLib, however, produces individually specified multipoint designs, which can become costly to synthesize and difficult to scale up when larger combinatorial libraries are required. htFuncLib extends this concept by using an Epistatic Neural Network (EpiNNet)^15^ to infer the compatibility of position-specific substitutions across multiple mutational backgrounds. Selected substitutions are then combined into experimentally accessible libraries,^16, 17^ which can be constructed economically using GGAssembler.^18^

FuncLib and htFuncLib enrich for folded, functional variants without directly identifying mutations that improve activity or selectivity toward specific substrates. Structure-based selectivity analysis can address this when structural models of the transition state are available^19^, but such information is often uncertain for new substrates, particularly when WT activity is weak or poorly selective.^20, 21^ A general strategy to direct htFuncLib sequence spaces toward target substrate activities without requiring solved transition-state complexes remains highly desirable.

Supervised machine learning can introduce this missing substrate-specific information. Data-driven methods have been used to guid directed evolution for more than two decades by learning experimentally measured sequence–function data and prioritizing variants for subsequent testing.^8, 22–25^ Their performance depends strongly on the size, diversity, and quality of the available training data, and many successful applications have relied on large datasets generated by high-throughput screening and sequencing.^23^ When activity and selectivity must be measured by methods like LC- and GC-MS, only a small fraction of a combinatorial library can be characterized. The initial library for obtaining training data is therefore critical in these low- to medium-throughput engineering campaigns. It must provide sufficient sequence diversity and fitness variation while containing enough functional variants to support model training.^10^ Structure-based design like htFuncLib offers a route to such an information-rich starting set. Experimental data from a compact htFuncLib library can then train a supervised model that redirects the sequence space toward the target function.

We developed a machine-learning-assisted enzyme-engineering (MLEE) workflow to connect stability-guided library design with specific function enrichment. First, htFuncLib defines a compact combinatorial sequence space enriched in multipoint mutants that are stable. A fraction of this library is then screened and sequenced with the target substrate. Second, supervised learning uses these sequence-function data to rank the amino acids available at each target position according to how beneficial or deleterious they are to the desired activity. The highest-ranked amino acids are combined into a compact second-round library enriched for the desired activity. We hypothesized that this workflow could redirect a substrate-independent htFuncLib sequence space toward substrate-specific function without exhaustive screening or a predefined productive substrate-binding pose. (**Figure 1**).

**Figure 1.**
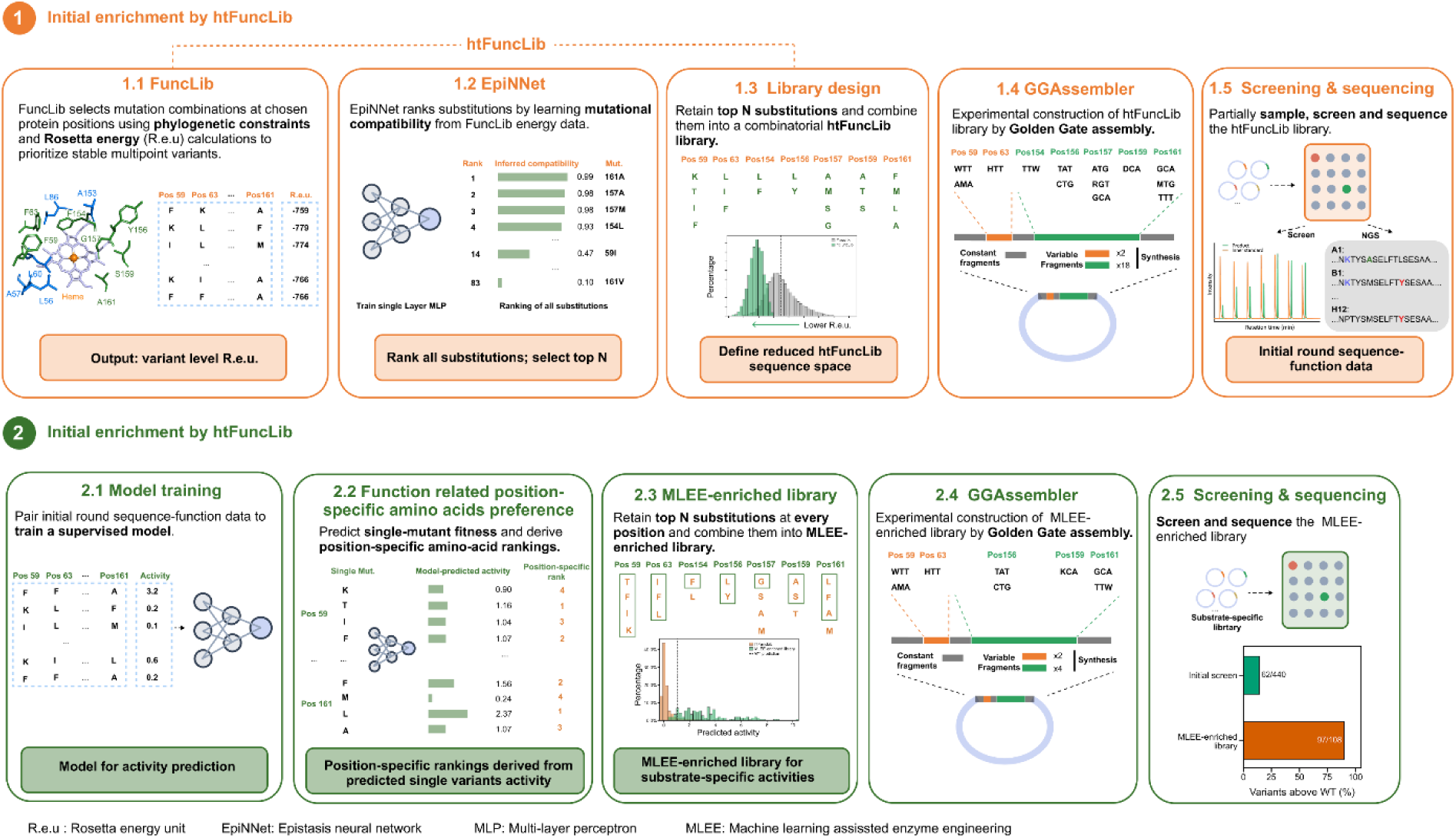
MLEE couples stability-guided library design with substrate-specific enrichment. htFuncLib first defines a compact sequence space enriched in substitutions that are compatible with one another but without directing to a specific reaction outcome. A fraction of this library is screened and sequenced, and a supervised model then ranks the allowed amino acids at each position by predicted activity. The highest-ranked amino acids define a second-round MLEE-enriched library.

Unspecific peroxygenases (UPOs) provide a relevant experimental system for evaluating this strategy. UPOs can catalyze oxyfunctionalization of a wide spectrum of substrates using hydrogen peroxide as the sole oxygen donor^26, 27^ and have been extensively engineered to improve expression, stability, activity, and chemo-, regio-, and stereoselectivity.^20, 28–31^ Their substrate scope extends from alkanes and aromatic compounds to structurally complex terpenoids.^28, 32–34^ UPO activity and selectivity depend on the collective architecture of the substrate-binding pocket, so optimization often requires coordinated mutations at several active-site positions.^20, 30^ Product formation and distribution also commonly require LC- or GC-MS analysis, limiting screening throughput and creating a strong need for compact libraries enriched in functional variants with the desired substrate conversion.

We first benchmarked the MLEE workflow on the near-complete four-position fitness landscapes of GB1^5^, TEV protease^35, 36^, and TrpB^37^. These datasets provide experimentally measured ground-truth data for evaluating both enrichment steps. Specifically, we tested if limited first-round screening data were sufficient to generate compact second-round libraries strongly enriched in high-performing variants. We then applied the workflow experimentally to *Mth*UPO-catalyzed β-damascone hydroxylation^12^, a fragrance-relevant terpenoid transformation^38^ that requires chromatographic analysis for activity and selectivity. The fraction of variants exceeding WT activity increased from 14% in the initial htFuncLib library to 90% in the MLEE-enriched library. The best variant, *Mth*UPO_11C5 (F59K/F63I/Y156L/S159A/A161F), showed an 11.8-fold increase with a turnover number of 1351 and greater than 99% regioisomeric excess (r.e.). toward 4-hydroxy-β-damascone. These results provide a proof-of-concept for MLEE as a strategy for transforming stability-guided htFuncLib sequence spaces into compact, substrate-specific libraries for enzyme engineering under constrained screening throughput.

## Results and discussion

### MLEE workflow as a two-step enrichment for specific function

The MLEE workflow consists of two major parts: 1) the generation of a high-throughput FuncLib (htFuncLib) library, followed by sequencing and testing a fraction of this library with a specific substrate; and 2) Machine-learning-derived activity prediction of htFuncLib designs to construct an enriched second-round library (**Figure 1**)

The first step of the MLEE workflow (**Figure 1**, upper panel) uses htFuncLib to define mutationally stable variants. This step does not require prior knowledge of the target substrate and does not explicitly optimize the target activity. Instead, it provides a multi-site library that is enriched in stable and foldable multipoint mutants relative to the full combinatorial sequence space.

This reduced htFuncLib library can still be too large to screen exhaustively. Therefore, the MLEE workflow samples a fraction of the initial htFuncLib library and uses the resulting sequence-function data to guide the second enrichment step toward substrate-specific activity (**Figure 1**, lower panel). A supervised activity-prediction model is trained from this sparsely sampled initial-round dataset and then applied to predict the fitness of the full htFuncLib library. The model is not used to directly select only a few top-predicted variants. Rather, the predicted activity ranking is used to identify position-specific amino-acid preferences associated with higher activity, which are then used to construct a second-round MLEE-enriched library biased toward the desired function.

### Benchmarking htFuncLib on complete mutational landscape as the first enrichment step

We evaluated MLEE using three experimentally characterized four-position fitness landscapes: GB1, the B1 immunoglobulin-binding domain of protein G;^5^ TEV protease, a sequence-specific cysteine protease that recognizes ENLYFQ↓S;^35, 36^ and TrpB, the β-subunit of tryptophan synthase^37^ (**Figure 2a**, **Table S1**). Each full 20⁴ sequence space contains 160000 variants, providing a stringent test of multi-site library enrichment. The three datasets also span distinct molecular functions and landscape structures, allowing the workflow to be assessed beyond a single protein system.

**Figure 2.**
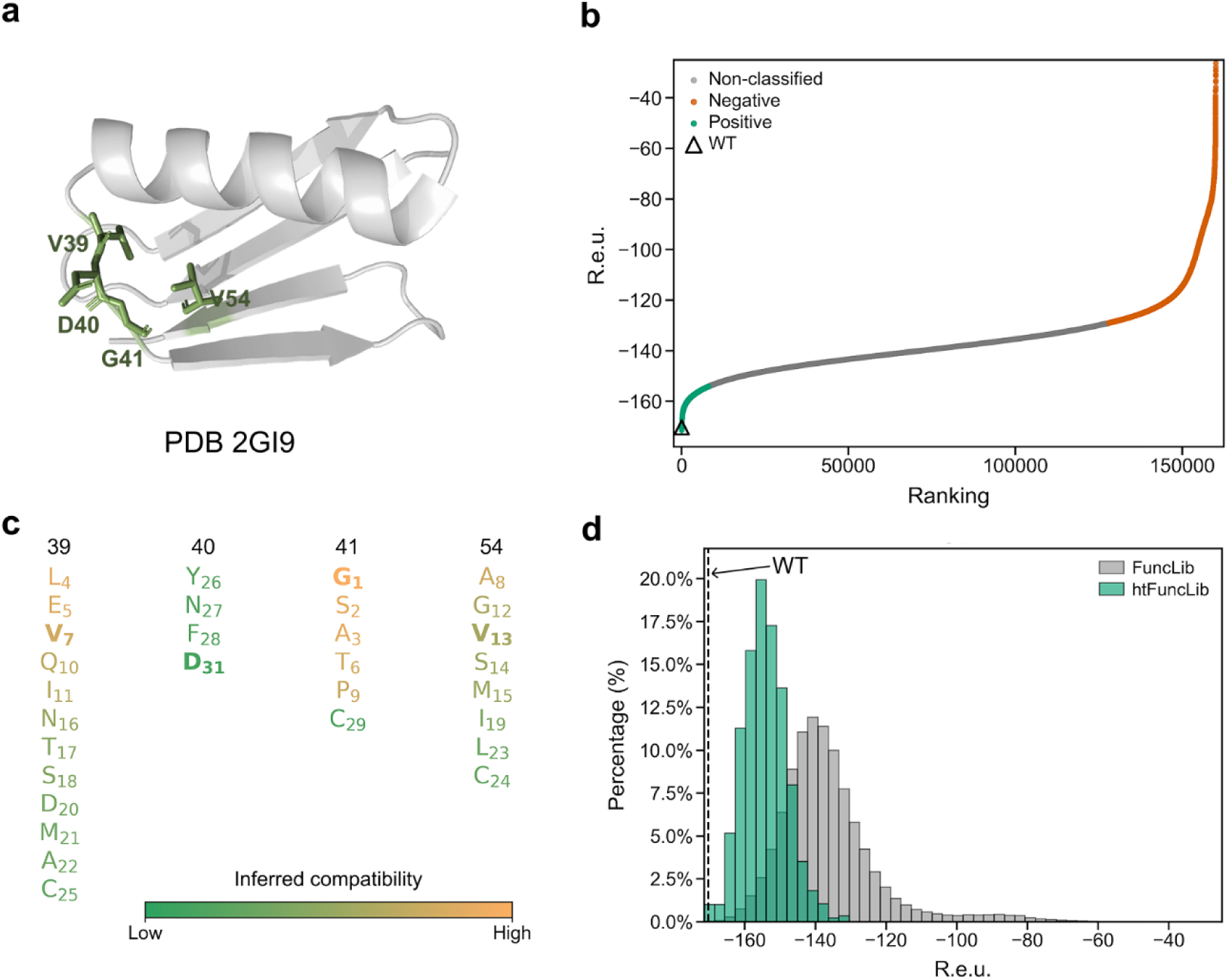
htFuncLib as the first enrichment step (GB1). **a. Structure of GB1**. Cartoon representation of GB1 based on PDB 2GI9, highlighting the four diversified positions, V39, D40, G41, and V54. **b. FuncLib energy distribution and classification of GB1**. Ranking of the complete 20^4^ GB1 sequence space, comprising 160,000 variants, by Rosetta energy (R.e.u.) calculated using FuncLib. Lower values indicate more favorable calculated energies. The lowest 5% of variants were labeled positive (green) and the highest 20% were labeled negative (orange) for EpiNNet training. 49 variants with Rosetta energy above −25 R.e.u. are cut for visual clarity but were retained in the ranking with negative class assignments. **c. Initial htFuncLib design for GB1**. Columns represent the four diversified positions, and amino-acid identities are ordered and colored according to their EpiNNet-inferred compatibility across mutational backgrounds. Subscripts indicate the global EpiNNet-derived rank among the 80 position-specific amino-acid identities, and WT residues are shown in bold. **d**. **Rosetta-energy enrichment for GB1**. The dashed line indicates the WT energy of -170.4 R.e.u. Corresponding analyses for TEV and TrpB are shown in **Figure S5**.

For each benchmark, we first calculated Rosetta energy scores for the complete 20⁴ combinatorial sequence space of each landscape using FuncLib (**Figure 1**, step 1.1, **Figure 2b**). In this step, standard FuncLib prefiltering based on MSA-derived amino-acid profiles and preliminary energy screening was omitted; instead, all 20 amino acids were retained at each of the four positions to match the experimentally characterized full landscapes. This exhaustive enumeration was computationally manageable for four positions, but it is not intended to represent the practical use case of htFuncLib. For larger multi-site designs, such as the UPO application described below, enumerating the full 20ⁿ sequence space would become computationally prohibitive and unnecessary, and the sequence space must instead be restricted by user-defined or multiple sequence alignment guided amino-acid sets before stability ranking and library construction.

The resulting energies were used to train EpiNNet, which ranked all position-specific substitutions in a single list, with each entry defined by both the amino acid and its target position. We applied a probability cutoff according to the available screening throughput, and all substitutions above this cutoff were retained, regrouped into amino-acid sets at each position, and combined to generate the htFuncLib library. (**Figure 1**, step 1.2 and 1.3). Known functional substitutions can also be incorporated at this stage when prior experimental information is available. For the benchmark simulation, we selected medium-throughput htFuncLib libraries of approximately 2200 variants for each landscape (**Figure 2c, Table S2**). These htFuncLib libraries strongly shifted toward energetically favorable variants compared to the full sequence space (**Figure 2d**). They, thus contained a substantially higher fraction of variants with fitness above the WT, increasing from 2.44% to 14.4% for GB1, from 0.21% to 2.38% for TEV, and from 0.69% to 8.06% for TrpB. (**Figure 3b**)

**Figure 3.**
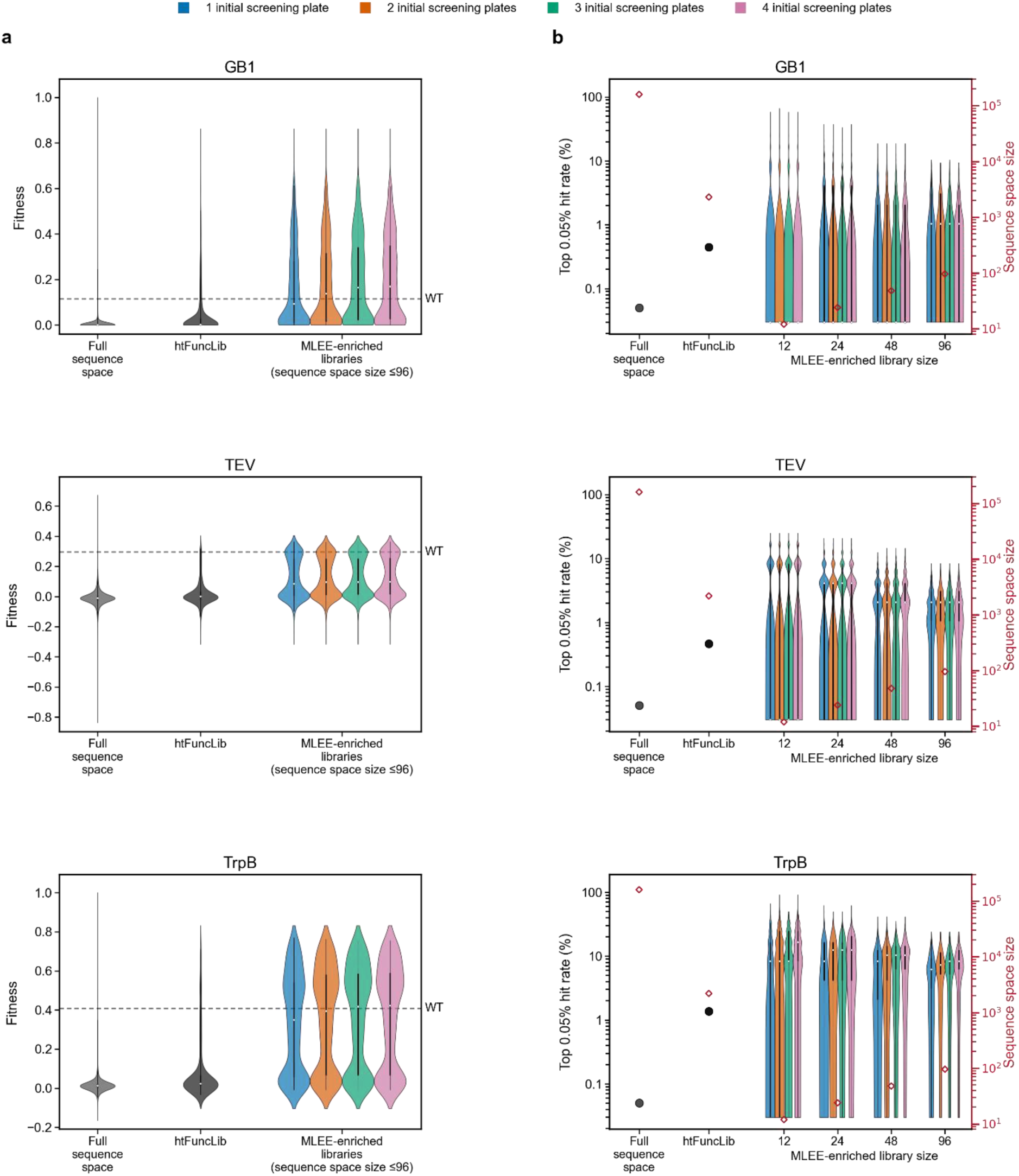
Two-step MLEE enrichment enables recovery of top-performing variants. **a. Fitness enrichment**. Fitness distributions of the full sequence space, htFuncLib, and all LEE-enriched libraries containing no more than 96 variants. At each position, amino acids were added sequentially from highest to lowest predicted single-mutant fitness, and all combinations of the retained amino acids were included in the resulting library. Violin plots summarize eligible libraries from 100 bootstrap simulations for each initial-screening condition. Fitness was normalized to the highest measured value in each benchmark landscape, and dashed gray lines mark WT fitness. **b. Top-variant recovery.** Top-0.05% hit rates for the full sequence space, htFuncLib, and exact 12-, 24-, 48-, and 96-variant MLEE-enriched libraries. Hit rate was calculated as the fraction of top-0.05% variants recovered from the complete library size. Only variants with measured fitness were used to define the top 0.05.

### Benchmarking Supervised learning on specific functions as the second enrichment step

From each selected htFuncLib library, we simulated initial-round screening by sampling one to four 96-well plates by using 100 bootstrapping simulations. The measured fitness values of the sampled variants from the corresponding dataset were paired with their sequences to train supervised regression models for activity prediction. Whereas EpiNNet used one-hot-encoded sequences to infer mutational compatibility from Rosetta energy calculation, the activity-prediction model used an AAindex-derived physicochemical representation (**Figure S6, Table S3,** see **AAindex_PC11 calculation** in **Supplementary Information**) to incorporate biochemical relationships among amino acids in the low- to medium-data regime.^39, 40^ The model optimization objective was explicitly tailored to activity prediction. Rather than seeking uniformly accurate predictions across the entire fitness range, we applied continuous fitness-dependent sample weights to emphasize the comparatively sparse high-fitness variants. These variants therefore contributed more strongly to the regression loss, directing model optimization toward high-function sequence regions relevant to the second-round library enrichment^41^

The trained activity-prediction model was then applied only to the full htFuncLib sequence space (**Figure 1**, step 2.1). Instead of directly selecting only the top-ranked predicted variants, we used the predicted ranking to identify which amino acids at each target position were preferentially associated with high predicted activity (**Figure 1**, step 2.2) Similar to EpiNNet, we predicted the WT-background single-variant fitness and used these values to rank the identities position by position. These predictions summarize residue preferences learned from the multipoint genetic backgrounds in the training data; they are not equivalent to direct measurements of isolated single mutants. Differences between predicted and measured single-mutant fitness were therefore examined as potential signatures of background dependence (Additional Results in the **Supporting Information**).^42, 43^

We computed MLEE-enriched libraries as sub-libraries of the original htFuncLib sequence space using the position-specific amino-acid rankings based on predicted single-mutant fitness. At each target position, amino acids were added progressively from the highest to the lowest rank. Different numbers of top-ranked amino acids could therefore be retained at different positions, for example, the top two amino acids at one position, the top three at another, and only the top-ranked amino acid at the remaining positions. (**Figure 1**, step 2.2) We enumerated all position-wise prefix combinations and retained libraries containing no more than 96 variants. This produced 906, 862, and 888 unique MLEE-enriched libraries for GB1, TEV, and TrpB, respectively. The MLEE-enriched libraries contained substantially more variants with fitness at or above WT than the starting htFuncLib libraries (**Figure 3a**). Across one to four initial screening plates, the median fraction ranged from 50.00% to 63.04% for GB1, 13.89% to 15.00% for TEV, and 45.83% to 54.76% for TrpB, compared to 14.4%, 2.38%, and 8.06% in the starting htFuncLib libraries, respectively (**Figure S7**). Thus, the supervised-learning step enriched functional variants beyond the initial stability-based enrichment.

We next tested whether the MLEE workflow could enrich globally top-performing variants, rather than only increasing the fraction of variants with fitness at or above WT. The top 0.05% of the measured landscapes comprised 75 GB1 variants, 76 TEV variants, and 80 TrpB variants. The starting htFuncLib libraries contained 10, 10, and 30 of these variants, corresponding to hit rates of 0.45%, 0.47%, and 1.37%. Thus, htFuncLib alone enriched for high functioning hits by 8.9-, 9.3-, and 27.2-fold above the 0.05% prevalence in the full GB1, TEV, and TrpB landscapes, respectively (**Figure 3b**).

Supervised learning increased this enrichment further. Substantial enrichment could even be achieved in 12-variant MLEE-enriched libraries. In 12-variant MLEE-enriched libraries, mean top-0.05% hit rates were 2.29-3.03% for GB1, 4.33-4.89% for TEV, and 11.41-16.86% for TrpB, depending on the initial screening size (**Figure 3b**). Increasing the second-round library to 96 variants improved recovery robustness: 61.3-66.8% of GB1, 84.0-87.2% of TEV, and 96.1-99.4% of TrpB libraries contained at least one top-0.05% variant. (**Figure S8**) When the initial screen was included, these probabilities rose to 70.7-89.0%, 89.5-96.9%, and 98.5-100%, respectively (**Figures S9**). Small MLEE-enriched libraries therefore maximized per-variant enrichment (**Figure 3b**), whereas larger libraries and the initial screen made improved recovery more reliable (**Figure S8**).

### MLEE enriches *Mth*UPO variants for β-damascone hydroxylation

We next tested the workflow experimentally by engineering *Mth*UPO for selective β-damascone oxyfunctionalization (Scheme 1).

**Scheme 1.**
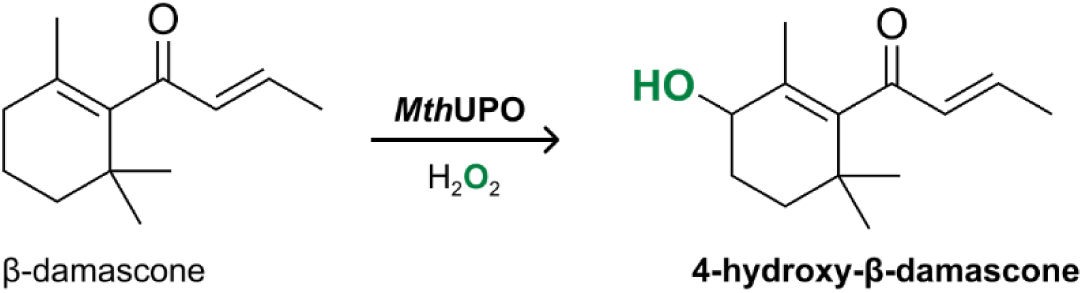
*Mth*UPO catalyzed the hydroxylation. *Mth*UPO uses H₂O₂ to convert β-damascone to 4-hydroxy-β-damascone.

### htFuncLib defines a compact *Mth*UPO active-site library

Twelve active-site residues, L56, A57, F59, L60, F63, L86, A153, F154, Y156, G157, S159, and A161, were selected for design. (**Figure 4a**) A sequence space of 529586 designs was calculated using FuncLib and evaluated using EpiNNet. Designs within the lowest 5% of Rosetta energy scores were labeled as positives, whereas designs scoring higher than the WT were labeled as negatives. To match the experimental screening capacity, we retained the 14 top-ranked mutations according to EpiNNet (**Table S5**) and additionally included F63I based on previous work.^12^ This selection produced an initial htFuncLib library of 2304 variants distributed across seven active-site positions (**Figure 4c, Table S6**). Relative to the initial FuncLib design pool, the htFuncLib library was strongly shifted toward energetically favorable designs, with 99.7% predicted to be more stable than the WT according to the Rosetta energy calculations. (**Figure 4d**) The library was then constructed by GGAssembler.

**Figure 4.**
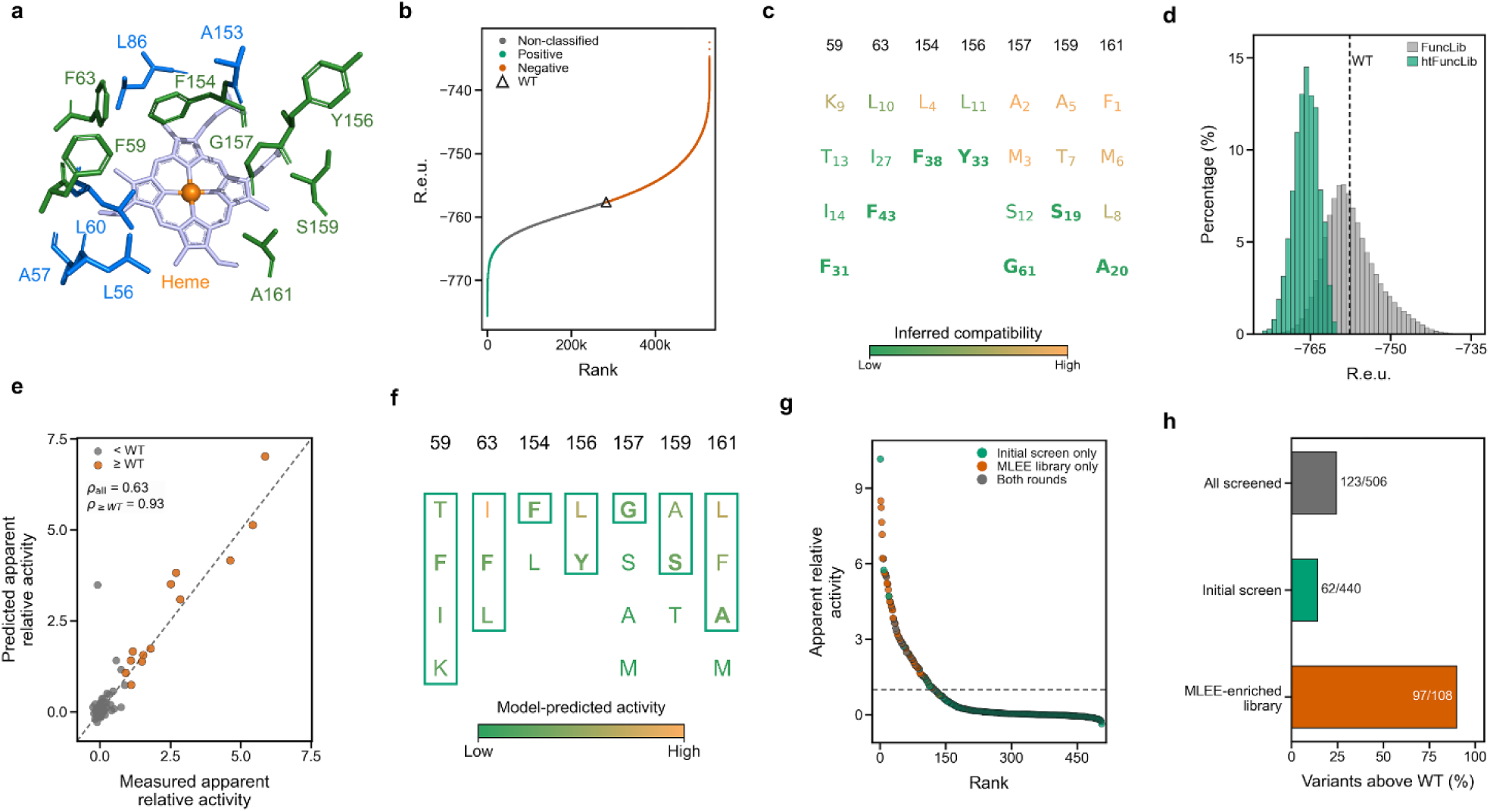
Application of MLEE to *Mth*UPO. a Structural selection of target positions. AlphaFold3^44^-predicted structure of *Mth*UPO showing the 12 active-site positions initially considered by FuncLib and the seven positions selected for htFuncLib construction (green). Heme shown in grey and iron in orange. **b**. **FuncLib energy classification**. Rosetta energy ranking of 529586 FuncLib designs. The lowest-energy (best) 5% of designs (green) and designs with energies above WT (orange) was used as positive and negative training data for EpiNNet, respectively. WT energy is marked by triangle. **c. Initial htFuncLib design**. EpiNNet ranked 82 candidate position-specific substitutions by their inferred compatibility across mutational backgrounds. The top 14 substitutions, the seven WT identities, and F63I from previous work^12^ were combined to generate a 2304-variant htFuncLib library. Colors indicate likelihood of stabilization, subscripts indicate global ranks, and WT residues are shown in bold. **d. Rosetta-energy enrichment**. Rosetta energy distributions of the FuncLib and htFuncLib designs. The dashed line indicates WT energy. **e. Model performance**. Predicted versus measured apparent relative activity for the held-out test set after training on data from 11 screening plates. Gray and orange points indicate variants below WT and at or above WT, respectively; ρ denotes the Spearman correlation coefficient. **f. Position-specific amino-acid ranking**. htFuncLib-accessible amino acids were ranked separately at each position using ML-predicted single-mutant activity. Amino acids for selected MLEE-enriched library are encircled. Colors indicate predicted activity, and WT residues are shown in bold. **g**. **Apparent relative activities of screened variants**. Apparent relative activities of all unique screened variants, ranked from highest to lowest and classified according to whether they were identified in the initial screen, the MLEE-enriched library, or both rounds. The dashed line indicates WT acticity. **h. Experimental activity enrichment**. Variants with activity above the corresponding WT accounted for 14% of initial htFuncLib variants (62/440), 90% of calibrated MLEE-enriched-library variants (97/108), and 24% of all unique screened variants (123/506).

### Initial screening yields a high-activity *Mth*UPO variant and training set

The experimental *Mth*UPO campaign involved a larger seven-position htFuncLib library than the benchmark datasets and a higher-dimensional sequence space for model training. We therefore continued first-round screening until a sufficiently informative dataset had been collected and a strong improved variant had been identified. After screening eleven 96-well plates, a five-point deign (F59K/F63I/Y156L/S159A/A161F), *Mth*UPO_11C5, was obtained, showing an apparent tenfold activity increase relative to the WT. At this point, the accumulated screening data were used for ML-guided library enrichment.

All screened transformants were sequenced, yielding 440 distinct first-round designs for subsequent ML analysis, among which 62 designs (14%) exceeded WT activity (**Figure 4h**, green and grey dots).

### Supervised learning and MLEE-enriched library test of *Mth*UPO

We next trained supervised activity-prediction models using the initial-round sequence-function dataset. Although the model showed only moderate ranking performance across all variants in the held-out test set (Spearman ρ_all_ = 0.63), its ranking performance was much stronger for variants with apparent relative activity above WT (Spearman ρ_≥WT_ = 0.93) (**Figure 4e**). This behavior was consistent with our intended use of the model: not to predict all activity with quantitative accuracy, but to enrich the next library toward high-activity sequence regions. We therefore predicted the WT-background single-variant activity for each htFuncLib-accessible amino acid and ranked the identities separately at each target position (**Figure 4f** and **Figure S4a**).

Using the predicted apparent relative activities, we selected a 144-variant MLEE-enriched library for second-round screening. (**Figure 4f, Table S8**) This library included 95 variants that had not been experimentally tested in the first round and was strongly biased toward high-predicted-activity regions of the initial htFuncLib sequence space, containing 9 of the top 10 and 74 of the top 100 predicted variants. The MLEE-enriched library was screened and sequenced using the same workflow as the initial htFuncLib library (**Figure 4g**, orange and gray dots).

Across both rounds, 506 distinct variants were screened and sequenced. In the initial htFuncLib screen, 62 of 440 variants (14%) exceeded WT activity. For the 108 MLEE-enriched-library variants, 97 (90%) exceeded WT activity (**Figure 4h**). The supervised-learning step therefore produced a 6.4-fold increase in the fraction of variants above WT. Importantly, improved activity was broadly distributed across multipoint variants rather than being confined to a small number of lightly mutated sequences. Among the measured second-round variants, 25 of 27 double mutants, 37 of 39 triple mutants, 25 of 27 quadruple mutants, and all six quintuple mutants exceeded WT activity, indicating that the MLEE-enriched library sampled a broad high-activity region rather than a single isolated optimum.

### Characterization of *Mth*UPO_11C5

*Mth*UPO_11C5 was purified for turnover number (TON) characterization under optimized reaction conditions with continuous H₂O₂ addition (**Figure S10**). The TON for formation of 4-hydroxy-β-damascone increased from 114 ± 6 for WT to 1351 ± 15 for *Mth*UPO_11C5 (**Figure 5a**), corresponding to an approximately 11.8-fold improvement under purified-enzyme conditions. As observed in our previous work, trace levels of 3-hydroxy-β-damascone and 4-oxo-β-damascone were detected under these reaction conditions for both WT and *Mth*UPO_11C5, but their abundance was too low for meaningful regioselectivity quantification. When comparing 4-hydroxy-β-damascone and 10-hydroxy-β-damascone, *Mth*UPO_11C5 reached >99% *r.e.* toward 4-hydroxy-β-damascone, compared with 39.8 ± 3.2% for WT (**Figure S11**). Notably, this high activity and regioselectivity were achieved without explicitly optimizing the initial library design for a β-damascone binding pose or a substrate-specific selectivity model.

**Figure 5.**
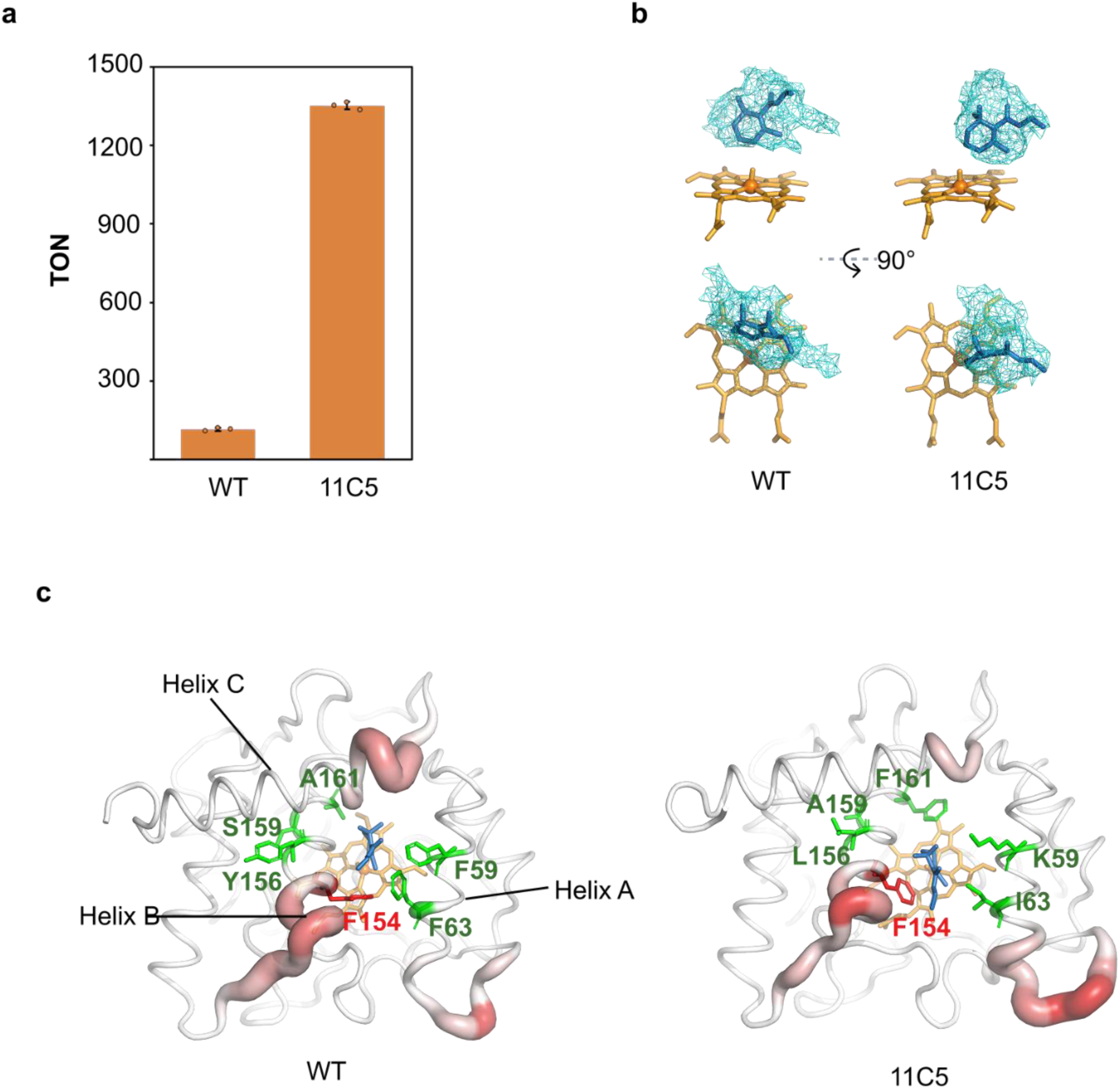
Characterization of *Mth*UPO_11C5 and residue-level activity effects. a. TON of purified enzymes in optimized condition. Purified enzymes were diluted with empty-vector culture supernatant to prepare 5× enzyme stocks and used at a final concentration of 50 nM. Reactions contained 1 mM β-damascone, 100 mM KPi buffer (pH 6.0), and 5% acetone. H₂O₂ was continuously supplied by syringe pump over 1 h to a final concentration of 1 mM, and products were extracted with EtOAc containing 250 μM (+)-valencene as internal standard. Data are from triplicate reactions. **b. β-damascone binding variance between WT and 11C5.** Representative MD snapshots showing the positioning of β-damascone when bound to compound I in the WT and 11C5. The mesh illustrates the conformational space sampled by the substrate in the simulations. **c MD-derived medoid structures of the WT and 11C5**. Residue flexibility is illustrated using a Putty representation (based on RMSF; see **Figure S14**).

Molecular dynamics simulations suggested that the improved activity and regioselectivity of *Mth*UPO_11C5 are associated with reorganization of the substrate-binding cavity. In simulations of the Compound I (Cpd I) state with bound β-damascone, the substrate adopted a more defined spatial arrangement in *Mth*UPO_11C5 than in WT (**Figure 5b**, **Figure S12**). This reduced conformational dispersion may increase the probability of productive alignment between the Fe(IV)=O of Cpd I and the C4 position of the β-damascone ring, thereby favoring hydrogen abstraction at the desired position.^20^ We also observed increased separation between helix A and helix B in *Mth*UPO_11C5 (**Figure 5c**, **Figure S13**), creating additional space to accommodate the substrate tail while positioning the β-damascone ring in a more favorable reactive orientation. This widened opening may also facilitate substrate access to the active site. Additional changes in local flexibility were observed near residues S200-N207 and A64S76 (**Figure S14**), indicating that the five active-site mutations affect both pocket geometry and protein dynamics.

## Conclusion

This work provides a proof-of-concept for MLEE as a two-step functional-enrichment strategy that connects the substrate-agnostic, stability-guided design of htFuncLib with substrate-specific enzyme engineering. Its central value lies at the library level: the model is not required to predict the activity of every sequence quantitatively or identify one exact optimum, but to increase the prevalence of favorable variants within an experimentally manageable second-round library. Benchmarking across GB1, TEV protease, and TrpB demonstrated that this strategy enriched globally high-performing variants of diverse fitness landscapes. In the experimental *Mth*UPO campaign, the fraction of variants exceeding WT activity increased from 14% in the initial htFuncLib library to 90% in the MLEE-enriched library, demonstrating that MLEE can enable multipoint enzyme engineering when functional screening is constrained by low- to medium-throughput analytical methods.

The unexpectedly high density of functional variants in the MLEE-enriched library suggests that the workflow exposed a locally smooth and functional region of the *Mth*UPO active-site landscape. Among the screened variants, from the MLEE-enriched library, improvement was broadly retained among variants carrying two to five active-site substitutions rather than being confined to one sequence or low-order mutants. This behavior reflects the complementary contributions of the two enrichment steps: native-state calculations first restricted diversification to structurally stable combinations, after which experimental activity data focused this sequence space toward β-damascone hydroxylation. A conceptually similar effect was reported for LAffAb, in which energy-guided mutation-tolerance mapping revealed a locally smooth multipoint landscape for antibody affinity maturation by limiting negative epistasis.^45^ The successful application of htFuncLib and ML to antibodies and an enzyme suggests the broad generality of this approach.

The final *Mth*UPO model used data from 11 screening plates, but the principal residue-selection decisions were already supported by the first four plates (Additional Results in the **Supporting Information**). 154F and 157G remained strongly favored over the alternative amino acids, leading to the same choices for the MLEE-enriched library. Residue-level library design may therefore remain robust with less initial data than required for uniformly accurate variant-level prediction. The necessary screening depth will still depend on library dimensionality, functional-variant density, experimental noise, and landscape complexity.

Future iterations could retain model-favored identities while adding designed mutations in selected active-site, second-shell, or distal positions. Cross-regional engineering of *Cvi*UPO has shown that such combinations can improve stability while maintaining or enhancing activity and selectivity.^46^ Iterative expansion would preserve the efficiency of focused library design while extending the accessible functional sequence space. MLEE thus provides a general framework for multi-site enzyme engineering when functional screening is constrained by analytical throughput.

## Materials and Methods

### Benchmark database

Benchmarking was performed using complete or near-complete four-site fitness landscapes^47^ for GB1^5^, TEV protease^35, 36^, and TrpB^37^. The analyzed positions were V39/D40/G41/V54 for GB1, T146/D148/H167/S170 for TEV, and V183/F184/V227/S228 for TrpB. Each theoretical sequence space contained 20⁴ = 160000 variants. Experimentally reported fitness values were used as ground truth and normalized to the maximum measured fitness in the corresponding landscape. The fitness values were normalized to the respective best variant. For TEV, any analysis requiring true activity used only variants supported by at least five sequencing reads. (**Table S1**)

### Chemicals and materials

See Supporting Information.

### Bacterial and yeast strains

*E. coli* DH10B (ThermoFisher Scientific, Waltham, US) was used for plasmid propagation. *S. cerevisiae* INV*Sc*1(ThermoFisher Scientific, Waltham, US) was used for enzyme expression and screening.

### htFuncLib library design

An AlphaFold3 model of *Mth*UPO truncated by 18 residues at the C-terminus was generated and used for all htFuncLib calculations. The protein sequence used for modeling is **Supporting Information**. Two rounds of FuncLib design were performed.^11^

In the initial round, diversification was introduced at positions L56, A57, F59, L60, F63, L86, A153, F154, Y156, G157, S159, and A161. C18, H88, and E158 were conserved, consistent with previous work^12^, and no amino acid identities were predefined at the target positions. Additional mutations reported previously^12, 20, 30^ were added to the second-round FuncLib design input (**Table S4**).

The second round used the adjacent-neighborhood strategy of the original htFuncLib workflow was applied.^17^ Ten adjacent neighborhoods defined by 6 Å sphere centered on the target positions (**Figure S15**), and one FuncLib calculation was performed for each neighborhood using amino-acid sets derived from the first round. The resulting designs were merged and deduplicated to give 529586 sequences with their Rosetta energy.

Designs were ranked in ascending order of Rosetta energy. The top 5% of designs were labeled as positive, whereas designs with Rosetta energies higher than that of the wild type were labeled as negative. (**Figure 4b**) These labeled sequences were one-hot encoded and used to train a MLP binary classifier, EpiNNet^15^. The sigmoid output of the classifier was interpreted as the estimated probability that a sequence belongs to the positive class. All 82 possible single substitutions at the target positions suggested by FuncLib were then evaluated *in silico* to estimate their stabilizing potential under the epistatic constraints captured by the trained model. Substitutions were ranked in descending order of activation value (**Table S5**).

Combinatorial libraries were constructed by sequentially introducing top-ranked substitutions into the wild-type background. Based on practical screening capacity, expansion of the first-round library was terminated at the 14th-ranked mutation, F59I. F63I was included based on previous work^12^, yielding the initial-round library design (**Figure 4c**, **Table S6**).

For the GB1, TEV, and TrpB, FuncLib was used to calculate Rosetta energy for the complete 20⁴ sequence space. The lowest 5% of designs were labeled as positives, and the highest 20% of designs were labeled as negatives. EpiNNet was then trained from these labels to rank all 80 position-specific amino acids in each benchmark landscape. The same one-hot EpiNNet architecture was trained on these labeled sequences and used to rank all 80 position-specific amino acids in one global list. Top 29 amino acids, and any missing WT amino acids of all positions were combined to define benchmark htFuncLib libraries. ( **Figure 2, Figure S5**)

### Machine-learning workflow for *Mth*UPO and benchmark simulations

The same general supervised-learning strategy was used for the experimental MthUPO data and the GB1/TEV/TrpB benchmark simulations. Sequence variants were encoded using AAindex_PC11 descriptors, which were generated by reducing AAindex-derived amino-acid physicochemical descriptors to 11 principal components, as described in the Supporting Information. For each multi-position variant, the AAindex_PC11 vectors of the amino acids at all target positions were concatenated to generate the input feature vector.

For model evaluation, data were split into training and test sets using an 80:20 split. The split was performed to preserve both the activity distribution and positional amino-acid representation between the training and test sets. A multilayer perceptron (MLP) regressor was optimized by grid search over network architecture, activation function, L2 regularization strength, sample-weighting exponent, and initial learning rate using 5-fold cross-validation on the training set, with mean squared error as the objective. The best model was then refit on the full training set and evaluated on the held-out test set. Optimization used the Adam optimizer with β₁ = 0.9 and β₂ = 0.999. The maximum number of training iterations was 500, and early stopping was applied. The hyperparameter grid is listed in **Table S15**.

Sample-weighted regression was used to emphasize variants with higher activity. Samples above the defined activity threshold were assigned larger weights according to their activity relative to the WT or threshold activity, controlled by a tunable weighting exponent. The weighting exponent was included in the hyperparameter grid.

For *Mth*UPO, models were trained using experimentally determined apparent relative activities from the initial htFuncLib screening round. The trained model was then applied to predict apparent relative activity across the full initial htFuncLib sequence space. For the GB1/TEV/TrpB benchmark simulations, models were trained from simulated initial screening subsets and then used to predict fitness across the corresponding htFuncLib-selected sequence space.

### Model-predicted single-mutant activity as a position-specific criterion for library enrichment

The trained model was used to predict fitness value (benchmark) and apparent relative activity (*Mth*UPO) of all possible single mutants allowed in the initial htFuncLib. At each position, amino acids were ranked from highest to lowest according to model-predicted values. Enriched combinatorial libraries were constructed using top-down selections. The Cartesian product of the retained residue set was defined the MLEE-enriched library.

### benchmark simulations

For each protein, the htFuncLib-selected library described above was used as the first-stage sequence space. Initial screening was simulated by sampling one, two, three, or four 96-well plates from the corresponding htFuncLib library. Because six wells per plate were reserved for controls, each plate corresponded to 90 screened variants, giving initial screening sizes of 90, 180, 270, and 360 variants. For each protein and screening size, 100 independent simulations were performed. Variants were sampled with replacement, duplicate variants were collapsed for model training, and the WT variant was included as a reference if it was not already sampled.

Candidate second-round MLEE-enriched libraries were generated as Cartesian products of top-ranked prefixes at the four positions, and all prefix combinations containing ≤96 variants were enumerated. Fitness summaries used only variants with eligible measured fitness, whereas hit-rate denominators used the theoretical library size. Top variants were defined as the highest-fitness 0.05% of measured sequences in each full landscape. When initial and second-round screening were analyzed together, duplicate sequences were removed and count as one.

### Library construction with GGAssembler

Initial and enrichment libraries were constructed using GGAssembler^18^ with pAGT572_Nemo 2.0, with minor adaptations in the wet-lab implementation. Briefly, GGAssembler designs Golden Gate assembly libraries by partitioning a target sequence into constant fragments without mutation sites and variable fragments with one or more target positions. These fragments can then be synthesized as oligonucleotides and assembled into the final library.

For each round, degenerate codons were first selected for all mutation positions (htFuncLib: **Table S7**; MLEE enriched library: **Table S9**). The *Mth*UPO coding sequence was then divided into five fragments: three constant fragments without mutation sites and two variable fragments. One variable fragment contained positions 59 and 63, and the other contained positions 154, 156, 157, 159, and 161. Oligonucleotides corresponding to the variable fragments were purchased from Eurofins Genomics (Ebersberg, Germany) (initial round: **Table S10**; enrichment round: **Table S11**).

Constant fragments were amplified using pAGT572_Nemo 2.0 carrying the wild-type MthUPO gene. Variable fragments were amplified from the purchased oligonucleotides. Primer sequences are listed in **Table S12**. All PCR products were purified before assembly.

Golden Gate assembly of the fragments, excluding the plasmid backbone, was performed as described previously.^48, 49^ The ligation product was then used directly as template for PCR amplification with primers F1_fwd and F5_rev (**Table S12**). The expected 992 bp product was verified by agarose gel electrophoresis, excised, and extracted using the NucleoSpin Gel and PCR Clean-up kit. The purified insert was subsequently assembled into pAGT572_Nemo 2.0 by Golden Gate assembly and transformed into *E. coli* DH10B to generate the library. Library composition was verified by Sanger sequencing (Eurofins Genomics, Ebersberg, Germany). The wild-type *Mth*UPO DNA sequence used for GGAssembler is provided in the **Supporting Information.**

### Enzyme screening

Transformation of the libraries into *S. cerevisiae* INVSc1 and enzyme expression in microtiter plates were carried out as described previously.^49^ Each plate contains three wells expressing WT enzymes, two wells containing empty vector, and one uninoculated well.

Screening reactions were performed in microtiter plates at a total volume of 500 μL. Each reaction contained 1 mM β-damascone, 1 mM H_2_O_2_, 100 mM KPi buffer (pH 7.0), 5% (v/v) acetone, and 100 µL culture supernatant added last. Reactions were incubated for 1 h, 30 ℃, and 300 rpm. After reaction completion, mixtures were extracted with 500 µL EtOAc containing 250 µM (+)-valencene as internal standard for 30min at 25 ℃ for 300 rpm. After phase separation by centrifugation at 1000 rpm for 10 min, the organic phase was analyzed by GC-MS on a Shimadzu GCMS-QP2010 Ultra instrument (Shimadzu, Kyoto, Japan) using helium as carrier gas.

Screening was performed using a multiple-injection in a single experimental run (MISER) GC-MS method.^49, 50^ Instrumental parameters are summarized in **Table S13**. Data were collected from 2 to 160 min in selected ion monitoring mode using *m/z* 189.00 for the internal standard and *m/z* 193.00 for the product. Relevant peaks were manually integrated. Apparent relative activity was calculated from the product/internal-standard signal and normalized to WT controls measured on the same plate. A representative chromatogram is shown in **Figure S16**.

Initial- and second-round activity measurements were placed on a common scale using 42 variants assayed in both rounds. An ordinary least squares regression was fitted to these shared measurements, and the resulting calibration equation was used to transform second-round activities onto the initial-round scale (**Figure S17**, y=0.22 + 0.87x, R^2^=0.68). The calibrated second-round data were then merged with the initial-round data. For variants measured in both rounds, the final activity was calculated as the weighted mean of the two round-specific values, using the number of screened transformants in each round as the corresponding weight.

### Multiplex NGS-based variant identification

After supernatants were collected for activity screening, cells from each well were inoculated onto SC-dropout (-uracil) agar plates according to the original microtiter plate layout and incubated at 30 °C for 24–48 h. Cells were lysed by transferring cell material into 96-well PCR plates containing 10 μL of 100 mM NaOH per well, followed by heating at 98 °C for 10 min. Lysates were neutralized by adding 50 μL of 22 mM Tris-HCl (pH 7.5). One microliter of neutralized lysate was used as template for PCR in a 10 μL reaction.

The first-stage PCR was performed using a touchdown program with annealing temperature decreasing from 65 °C by 0.7 °C per cycle for 14 cycles. Primers NGS_inner_fwd and NGS_inner_rev were used at 0.02 μM each. After the first PCR stage, 2 μL of barcode primer mix was added to the corresponding well position to give a final primer concentration of 0.2 μM. (**Table S14**) The second-stage PCR was then performed with annealing at 66 °C for 30 cycles.

After amplification, 5 μL PCR product from each well of a plate was pooled into 120 μL of 100 mM EDTA solution at pH 8.0 to quench the reaction, giving a final EDTA concentration of 20 mM. The pooled PCR product from each plate was separated by agarose gel electrophoresis, and the expected 510 bp band was excised and purified by gel extraction. Each plate was submitted as one sample for NGSelect Amplicon MiSeq sequencing at Eurofins Genomics.

For each plate, sequencing data were received as paired forward and reverse FASTQ.gz files. Data processing was adapted from evSeq^51^ as detailed in Supplementary information.

### Protein expression, purification, and TON determination

Enzyme expression for screening and turnover number (TON) measurements was performed in microtiter plates. For purification, material from four plates each of WT and 11C5 was pooled separately. For preparative synthesis of 4-hydroxy-β-damascone, 11C5 was expressed in shake flasks and subsequently concentrated. All procedures were performed as described previously^52^.

Enzyme concentration was determined by CO difference spectroscopy. TON measurements were performed in glass vials. For batch reactions, a master mix was prepared to give final concentrations of 1 mM substrate, 5% acetone, 100 mM KPi buffer, and H_2_O_2_ in a total volume of 500 μL. Unless otherwise stated, reactions were performed at pH 7.0; for pH optimization, pH 6.0, 7.0, and 8.0 were tested.

For reactions with continuous H_2_O_2_ addition, 400 μL of master mix without H_2_O_2_ was first prepared, followed by addition of 100 μL of 5 mM H_2_O_2_ over 1 h using a syringe pump. Reactions were performed at 30 °C with continuous shaking. Products were extracted by addition of 500 μL EtOAc containing 250 μM (+)-valencene, followed by shaking at 25 °C. The organic phase was analyzed by GC-MS under the conditions listed in **Table S13**. Samples were analyzed in triplicate unless stated otherwise; troubleshooting experiments were performed in duplicate. Quantification was based on a calibration curve generated with 4-hydroxy-β-damascone (**Figure S110**; details in the Supporting Information). TON was defined as the amount of product formed per amount of enzyme after 1 h and calculated according to calibration curve (see **Supplementary Information**, **Figure S18**).

### MD simulation

Initial structures of the wild-type (WT) and 11C5 MthUPO variants, including the heme cofactor, Mg^2+^ ion, and bound β-damascone, were generated with Chai-1,an open-access AlphaFold3-based platform for biomolecular interaction prediction. Five models were selected for each enzyme-substrate complex and used as starting structures for molecular dynamics (MD) simulations,^53^ an open-access AlphaFold3-based platform for biomolecular interaction prediction. Five models were selected for each enzyme-substrate complex and used as starting structures for molecular dynamics (MD) simulations.

Each system was solvated in an 8.0 × 8.0 × 8.0 nm^3^ box containing approximately 16,000 water molecules. Systems were neutralized with 12 K^+^ ions, and additional KCl was added to a final concentration of 100 mM. Energy minimization was performed in 50 cycles, each consisting of 100 steepest-descent steps followed by one conjugate-gradient step. The minimized systems were equilibrated with a 200 ps NVT simulation using positional restraints of 1000 kJ mol^-1^ nm^-2^ on heavy atoms. This was followed by three NPT equilibration steps with gradually reduced positional restraints, from 1000 to 100 kJ mol^-1^ nm^-2^, over a total of 1 ns. Production simulations were then run for 500 ns in the NPT ensemble without positional restraints using a 2 fs time step. Coordinates were saved every 10 ps.

To mimic a near-attack conformation of β-damascone in the active site, a harmonic distance restraint was applied between the C4 atom of β-damascone and the activated oxygen of compound I (CpdI), with a target distance of 2.7 Å and a force constant of 10,000 kJ mol^-1^ Å^-2^, during both equilibration and production runs.

All MD simulations were performed with GROMACS 2021.4.^54^ The AMBER14SB^55^ force field and TIP3P water model^56^ were used. Parameters for CpdI and the axial cysteine were taken from the literature.^57^ The geometry of β-damascone was optimized with Gaussian 16,^58^ at the TPSSTPSS/def2-TZVP level, and the resulting structure was subsequently used to compute RESP^59^ atomic point charges at the same level of theory. The ligand was parameterized with the antechamber module in AMBER 23^60^, using GAFF2^61^, and converted to GROMACS format with ACPYPE^62^.

van der Waals interactions were switched off between 1.1 and 1.2 nm. Long-range electrostatic interactions were treated with the particle mesh Ewald method^63, 64^ using fourth-order interpolation, a real-space cutoff of 1.2 nm, and optimized fast Fourier transform parameters with a grid spacing of approximately 0.16 nm. Bonds involving hydrogen atoms were constrained using the fourth-order double-iteration parallel linear constraint solver (P-LINCS).^65^ Verlet neighbor lists were updated every 20 fs, with the neighbor-list radius determined automatically. Simulations were carried out in the NPT ensemble with isotropic Parrinello-Rahman pressure coupling (1 bar, τ_p_ = 2 ps, κ_p_ = 4.5 × 10^-5^ bar^-1^)^66^, and Bussi-Donadio-Parrinello velocity-rescaling thermostatting (300 K, τ_T_ = 0.2 ps)^67^. Molecular structures were visualized with PyMOL (Schrödinger, LLC).

## ASSOCIATED CONTENT

### Supporting Information

The following files are available free of charge

All screened and sequenced deigns: Supplementary Table 1.xlsx

Original data, MD data, code for benchmark and ML will be available once the paper is accepted.

## AUTHOR CONTRIBUTIONS

Conceptualization: L.W., S.J.F., and M.J.W.; Wet lab: L.W., and L.F., machine learning: L.W; molecular dynamic simulation: M.B.H.; data curation and visualization: L.W., and M.B.H.; draft: L.W., M.H.B., and L.F. Review and editing: L.W., M.B.H., S.J.F., and M.J.W.; Supervision: S.J.F., and M.J.W..

