## Supporting Information for "Combining Stability-Centered Atomistic Design with Machine Learning for Targeted Enzyme Optimization"

**Chemicals and materials.**

β-damascone (≥90 %), (+)-valencene (>70 %), and $H_2O_2$ (30 % (w/w) in $H_2O$) were purchased from Sigma-Aldrich (St. Louis, US)

Acetone (Rotisolv ≥99.9 % GC Ultra Grade), and EtOAc (Rotisolv ≥99.9 % GC Ultra Grade) were purchased from Carl Roth (Karlsruhe, Germany).

D-Galactose, Peptone and Synthetic Complete Mixture (Kaiser) Drop-Out (-uracil) for *S. cerevisiae* culture were purchased from Formedium (Hunstanton, GB). LB medium and LB-agar (Luria/Miller) powder for *E. coli* culture were purchased from Carl Roth (Karlsruhe, Germany).

BsaI was purchased from New England Biolabs (Ipswich, US). T4 DNA Ligase from Promega (Madison, US). All PCR was carried out with Phusion™ Plus DNA Polymerase from Thermo Fisher Scientific (Vilnius, LT).

GC column (OPTIMA 5 MS, 30 m L, 0.25 mm ID), and Gel and PCR Clean-up kit (NucleoSpin) was purchased from MACHEREY-NAGEL (Düren, DE).

***Mth*UPO sequence used for AF3**

AGFDTWSPPGPYDVRAPCPMLNTLANHGFLPHDGKDITREQTENALFEALHINKTLA
SFLFDFALTTNPKNTSTFSLNDLGNHNILEHDASLSRADAYFGNVLQFNQTVFDETKT
YWEGDTIDLRMAAKARLGRIKTSQATNPTYSMSELGDAFTYGESAAYVVVLGDKES
RTVKRSWVEWFFEHEQLPQHLGWKRPAASFEEEDLNSSMEEIEKYTKELEGSNSTS*G*
*SQKHRRRLPRRRAHFGF(Truncated)*

**WT *Mth*UPO DNA sequence used for GGAssembler**

signal peptide from α Galactosidase (*S. cerevisiae*), and *TwinStrep-GFP11 tag*

ATGTTTGCTTTTTATTTCTTGACTGCTTGTATTTCTTTGAAAGGTGTTTTTGGAGCA
GGTTTTGACACTTGGTCACCACCTGGACCCTATGATGTTAGGGCTCCTTGTCCGAT
GTTGAATACATTGGCTAATCATGGTTTCTTACCACATGATGGCAAAGATATTACTC
GTGAACAGACAGAAAACGCCTTGTTCGAAGCATTGCACATCAACAAAACCTTAG
CCAGCTTTCTGTTTGACTTTGCATTAACAACGAATCCGAAGAATACCTCGACGTTT
TCACTGAACGACTTAGGCAATCACAACATTTTGGAACATGATGCATCACTAAGTA
GGGCTGATGCGTACTTTGGGAATGTTCTACAGTTCAATCAAACTGTCTTTGATGA
GACTAAAACCTACTGGGAAGGAGATACTATTGATTTGAGAATGGCAGCCAAAGC
TAGACTAGGTAGAATCAAGACATCTCAAGCTACTAATCCAACGTATTCCATGTCG
GAATTAGGAGATGCTTTCACATATGGGGAATCTGCTGCGTATGTGGTAGTGTTAG
GTGACAAAGAGTCTCGTACTGTCAAAAGATCCTGGGTTGAATGGTTCTTCGAACA
TGAGCAACTTCCTCAACATCTTGGTTGGAAAAGACCAGCAGCATCCTTCGAAGAA
GAAGATCTGAACTCATCAATGGAGGAGATTGAGAAGTACACCAAGGAATTGGAA
GGTAGCAACTCTACAAGTGGTAGTCAAAAGCATAGAAGGAGACTTCCAAGAAGA
AGAGCTCACTTTGGCTTT*TCGGGTGGTTCTGCTTGGTCACATCCACAATTTGAAAAAG*
*GTGGAGGTTCAGGTGGAGGTTCGGGTGGTTCTGCTTGGTCACATCCACAATTTGAAAA*
*AGATGGTGGTTCTGGTGGTGGTTCTACTAGTCGTGATCATATGGTTCTTCATGAATATGT*
*TAATGCTGCTGGTATTACTTGA*

**AAindex_PC11 calculation**

AAindex_PC11 descriptors were calculated from AAindex-derived amino-acid physicochemical descriptors. A 20 × 566 amino-acid descriptor matrix was obtained from the AAindex data source (https://github.com/amckenna41/aaindex.git)[1]. Each descriptor was standardized to have mean 0 and standard deviation 1 across the 20 canonical amino acids. Principal component analysis was then applied, and the first 11 principal components were retained. The cumulative explained variance is shown in **Figure S5**, and the resulting AAindex_PC11 descriptor table is provided in **Table S3**.

**Calibration curve for 4-hydroxy-β-damascone**

Seven stock solutions of 4-hydroxy-β-damascone in acetone were prepared by serial dilution from a 50 mM stock solution to final stock concentrations of 0, 0.1, 0.2, 0.4, 1, 2, and 4 mM. For each calibration sample, 25 µL of stock solution was added to 475 µL assay solution to give final 4-hydroxy-β-damascone concentrations from 0 to 200 µM, with 5% acetone and 100 mM KPi buffer at pH 7.0 in a total volume of 500 µL. All calibration samples were prepared in triplicate, extracted, and analyzed as described for TON determination. The calibration curve is shown in **Figure S18**.

**NGS data processing**

For each plate, sequencing data were received as paired forward and reverse FASTQ.gz files. Data processing was adapted from evSeq.[2]

Forward and reverse reads were first paired according to flow-cell coordinates in the read headers, and unpaired reads were discarded. Read pairs were subjected to quality control using a minimum length of 270 bases and a minimum Phred score of 25. A read pair was retained when at least one read passed quality control; read pairs were discarded only when neither read passed quality control. Read pairs were assigned to wells only when both forward and reverse barcodes fully matched a barcode pair assigned to a well in that plate. Unassigned read pairs were discarded.

Reads passing barcode assignment were aligned to the reference amplicon sequence using semiglobal alignment with match score 1.0, mismatch score 0, gap opening penalty -2.0, gap extension penalty -0.5, and no end-gap penalty. Alignments were rejected if they contained any insertion in the read or more than 15 mismatches for initial round and 12 for MLEE-enriched library.

Composite base and amino-acid alignments were constructed for each well. For base-level composite alignments, base calls from forward and reverse reads were combined using Phred quality scores and base counts. When both reads gave the same base at a position, the base was called with count 2 if both Phred scores were ≥30. If only one read passed the Phred 30 threshold, the base was called with count 1 from the passing read. If both reads were below the Phred 30 threshold, the base with the higher Phred score was used with count 1. When the two reads disagreed, the base with the higher Phred score was used with count 1. If only one read covered the position, that read was used with count 1. Gap calls were treated separately. Counts for each base and gap were summed at each position for each well.

For amino-acid composite alignments, each codon was translated after considering base quality and gaps. Codons containing a gap were assigned to the gap channel. Codons containing any base with Phred score <30 were excluded from amino-acid counting for that read pair. Codons with all bases having Phred score ≥30 were translated to the corresponding amino acid or stop codon. Amino-acid, gap, unknown, and stop-codon counts were summed at each position for each well.

Variable positions were identified from gap-excluded base and amino-acid frequencies using a normalized L1 distance from the reference state. This distance is equivalent to one minus the reference frequency for a one-hot reference state. A position was called variable when the distance was >0.2, corresponding to reference base or amino-acid frequency <0.8. The first and last three DNA positions and the first and last amino-acid positions were excluded from variable-position calling. Wells were flagged as failing the mutation-count filter when more than 15 DNA-level variable positions were detected. The mutation-count filter was applied to the number of variable DNA positions; amino-acid variable counts were recorded separately.

Haplotype calling required at least 10 good read pairs. Final mutation calls for each well were assigned from the most frequent amino-acid haplotype across all variable positions, corresponding to the most abundant mutation combination among read pairs in that well. Position-wise base and amino-acid count tables were also retained as supporting information for each well.

**Reference sequence for NGS data processing**

AACGCCTTGTTCGAAGCATTGCACATCAACAAAACCTTAGCCAGCTTTCTGTTTG
ACTTTGCATTAACAACGAATCCGAAGAATACCTCGACGTTTTCACTGAACGACTT
AGGCAATCACAACATTTTGGAACATGATGCATCACTAAGTAGGGCTGATGCGTAC
TTTGGGAATGTTCTACAGTTCAATCAAACTGTCTTTGATGAGACTAAAACCTACTG
GGAAGGAGATACTATTGATTTGAGAATGGCAGCCAAAGCTAGACTAGGTAGAAT
CAAGACATCTCAAGCTACTAATCCAACGTATTCCATGTCGGAATTAGGAGATGCT
TTCACATATGGGGAATCTGCTGCGTATGTGGTAGTGTTAGGTGACAAAGAGTCTC
GTACTGTCAAA

**Enzymatic preparation of 4-hydroxy-β-damascone.**

The reaction was carried out in a round-bottom glass bottle with magnetic stirring and water bath at 30 °C. The reaction mix was in total 80 mL with 4 mM β-damascone, 5% acetone, 100 mM pH 7.0 Kpi, 1 mM $H_2O_2$ at the beginning and 5 mL of 5 mM $H_2O_2$ addition in the reaction period of 7.5 h, and 500 nM of crude 11C5 enzyme expressed and concentrated from yeast expression in shake flasks. The reaction mixture was extracted 3 times with EtOAc, washed with brine and then with $Na_2SO_4$. And then the extractant was concentrated by rotary evaporation and subjected to purification

The crude extract from the biotransformation was purified by flash column chromatography ($SiO_2$, pentane : $Et_2O$ 5:1 → 1:1). Analytical thin-layer chromatography (TLC, Macherey-Nagel

250) was performed to observe the product containing fractions, visualized by staining with a Cer solution. Purification through column chromatography was performed with silica gel (Macherey-Nagel, 40-63 μm).

NMR spectroscopy of $^{1}$H-, $^{13}$C- and 2D-NMR experiments were performed on Bruker AVANCE NEO (νL(1H)=600 MHz) equipped with DUL cryoprobe (**Figure S19**. The probe is equipped with a z-Gradient coil. Data analysis was performed using the Mestrelab Mestrenova software. The residual solvent signal of the deuterated solvents was used to calibrate the chemical shift scale of the NMR spectra. Chemical shifts δ are given in ppm, J coupling constants are given in Hz and were determined manually. The abbreviations used for multiplicities are s (singlet), d (doublet), t (triplet), q (quartet) and m (multiplet). Structure elucidations of novel compounds were supported by the usage of 2D-NMR experiments (namely: $^{1}$H-$^{13}$C HSQC, $^{1}$H-$^{13}$C HMBC and $^{1}$H-$^{1}$H COSY). GC/MS analyses were carried out with an Agilent 7890B GC with 5977B GC/MSD and Gerstel MPS Robotic XL with KAS 4C injector (**Figure S20)**. Samples were analysed on an Optima 5HT column, 30 m x 250 μm i.d. x film thickness 0.25 μm). Carrier gas, He; injector temp.: 60 °C to 300 °C at 12 °C/min, splitless; temp. program: 50 °C (isothermal 1 min) to 300 °C, at 20 °C/min and held isothermal for 6.5 min at 300°C; FID: 300 °C, $H_2$: 30 mL/min, $N_2$: 25 mL/min, MSD: ion source: EI 70 eV, 230 °C; detector: quadrupole, EI mass spectra were acquired over the mass range of 30 – 650 amu.

**$^{1}$H NMR** (600 MHz, $C_6D_6$) δ 6.59 (dq, *J* = 15.7, 6.9 Hz, 1H), 6.09 (dq, *J* = 15.7, 1.6 Hz, 1H), 3.65 (t, *J* = 4.8 Hz, 1H), 1.78 – 1.68 (m, 1H), 1.61 (s, 3H), 1.58 – 1.53 (m, 2H), 1.36 (dd, *J* = 6.9, 1.7 Hz, 3H), 1.27 – 1.18 (m, 1H), 1.10 (s, 3H), 1.09 (s, 3H) ppm.

**$^{13}$C NMR** (151 MHz, $C_6D_6$) δ 199.4, 144.8, 143.5, 134.5, 131.3, 68.8, 35.0, 34.2, 29.0, 27.9, 18.1, 17.9 ppm.

### Additional results

#### Comparison between measured single mutant fitness and model-predicted single mutant fitness for position-specific ranking of amino acids for specific functions.

We compared MLEE-enriched libraries generated using position-specific amino acid rankings based on either measured or model-predicted single mutant fitness. Across all libraries with sequence-space no greater than 96, the measured ranking produced higher median fractions of variants with fitness at or above WT for GB1 (71.9% versus 50.0–63.0%) and TrpB (58.1% versus 45.8–54.8%). Model-predicted ranking was consistently better for TEV, where the median increased from 8.45% to 13.9–15.0%. For globally top variants, model ranking increased the mean top-0.05% hit rate from 1.32% to 1.87–2.24% for GB1 and from 1.87% to 3.06–3.29% for TEV. Measured ranking remained better for TrpB (15.9% versus 8.98–11.9%; **Figure S1**).

The upper ends of the distributions revealed an additional advantage of model-based ranking. At a fixed library size, the maximum obtained using the measured ranking represents the best library accessible under that ordering. Model ranking generated libraries beyond this measured-ranking limit. For a 12-variant GB1 library, the maximum top-0.05% hit rate increased from 8.33% to 66.7%. For TEV, the maximum fraction of variants with fitness at or above WT increased from 50.0% to 83.3%, and the maximum top-0.05% hit rate increased from 16.7% to 25.0%. A smaller advantage was also observed for TrpB, where the maximum hit rate of a 48-variant library increased from 31.2% to 41.7%. These upper-tail gains remained when only the ten models with the highest held-out test-set Spearman correlations were analyzed: the corresponding 12-variant maxima were 50.0% versus 8.33% for the GB1 top-0.05% hit rate and 75.0% versus 50.0% for the TEV fraction at or above WT (**Figure S2**). Thus, the gains were not restricted to poorly performing models. Model-predicted single mutant fitness was not universally superior, but it could promote or demote residues according to patterns learned from multi mutant backgrounds, thereby enabling compact high-function libraries that were inaccessible under the measured single mutant ordering at the same screening capacity.

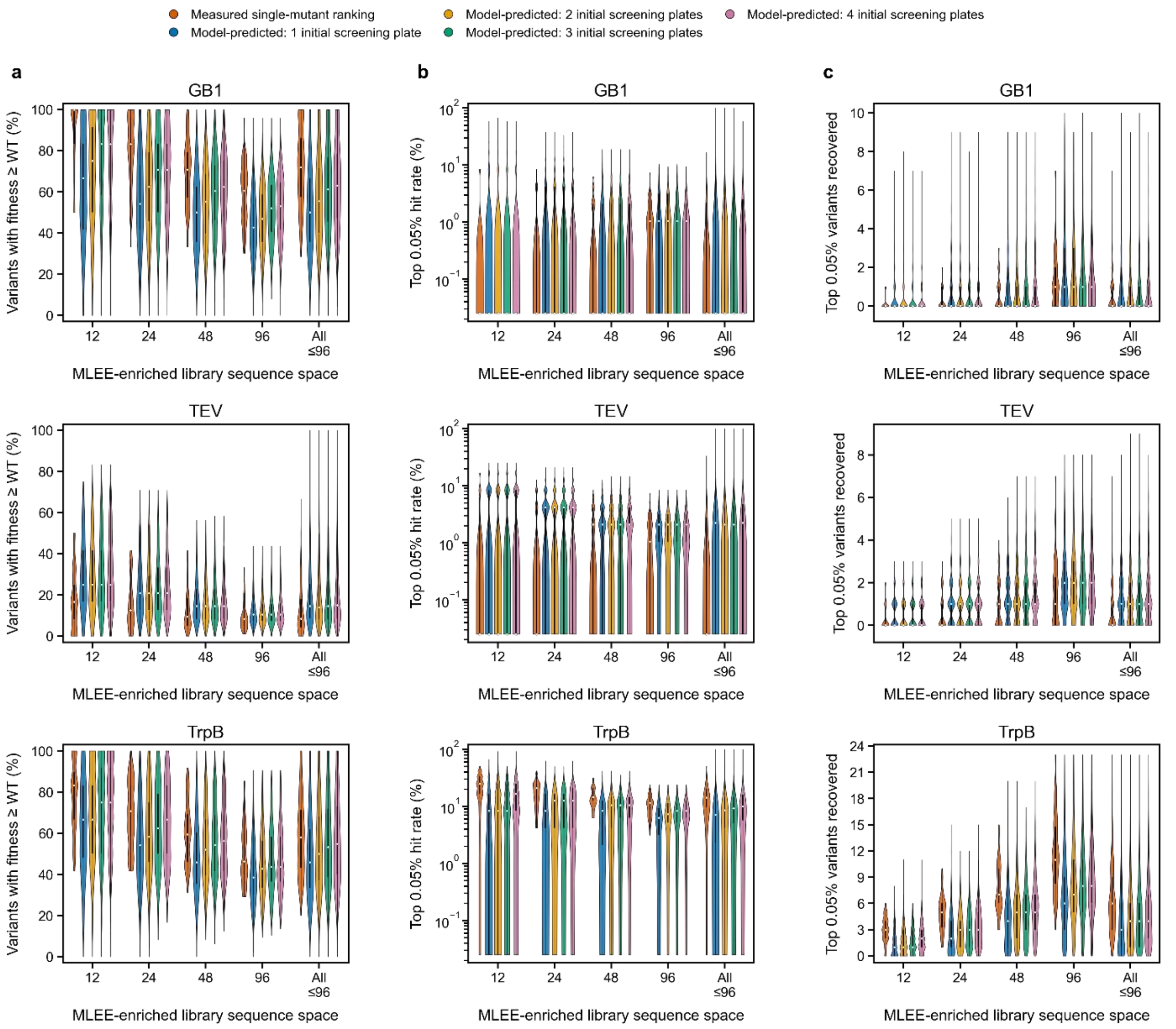


**Figure S1**. **Measured versus model-predicted single mutant ranking across all simulations.** Violin plots compare MLEE-enriched libraries with exact sizes of 12, 24, 48, and 96 variants and all attainable libraries with size ≤96**. a. variants with fitness at or above WT**, **b. top-0.05% hit rate**, and **c.top-0.05% variants recovered**. Model-derived distributions include 100 simulations per initial-screening condition. TEV fitness measurements required at least five reads. Internal black lines show the interquartile range, thin lines show adjacent values, and white points show medians.

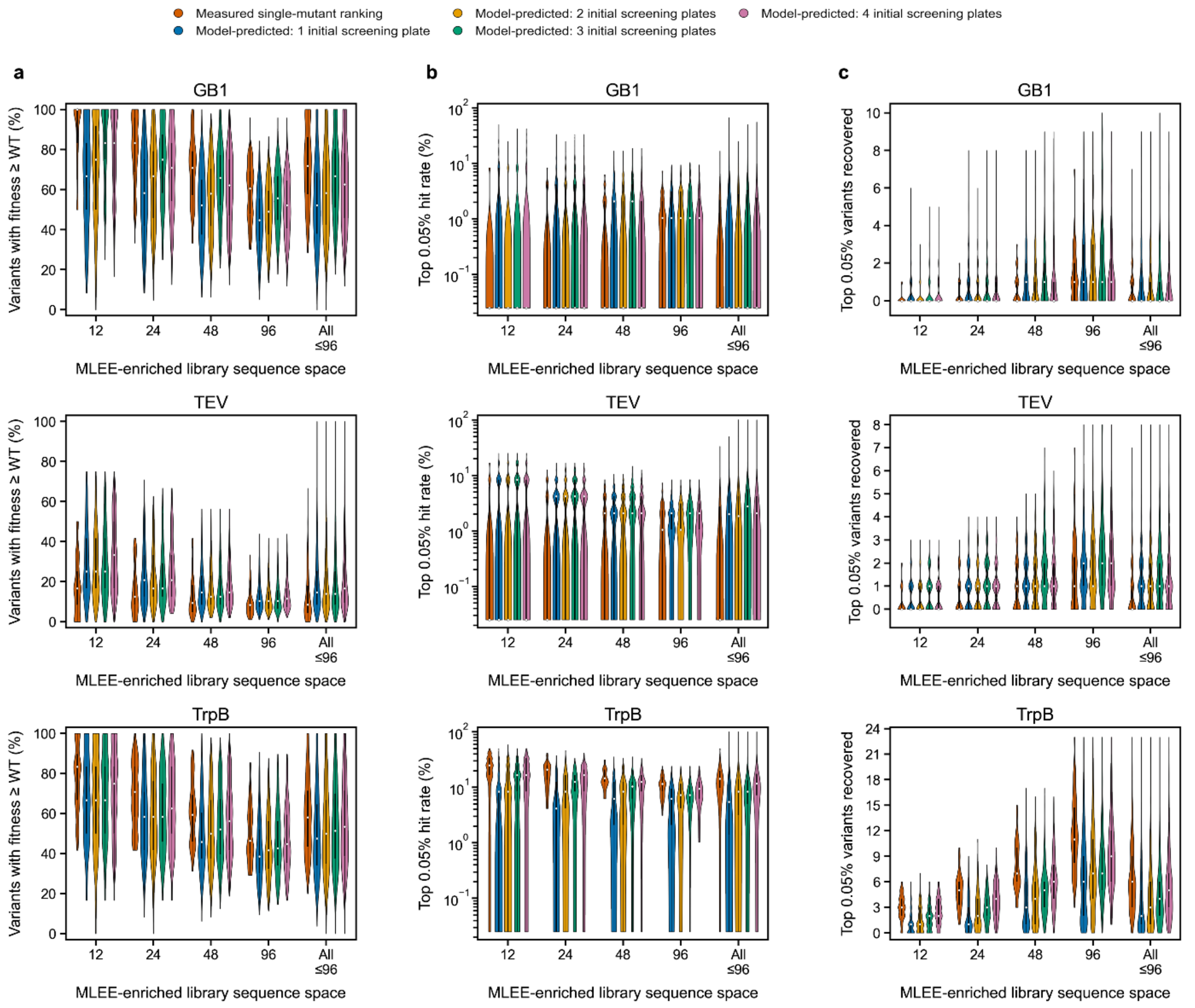


**Figure S1**. **Ranking comparison for models with the highest held-out performance.** The analysis was repeated using the ten simulations with the highest held-out test-set Spearman correlation for each protein and initial-screening condition. The measured-ranking reference and all metric definitions are identical to **Figure S1.**

**Four initial screened plates reproduced the MLEE-enriched library selected using 11 plates**

To determine whether fewer initial measurements were sufficient for MLEE library design, we trained the regression model using only data from the first four screened plates. Though not as good as the model trained from all 11 plates, the resulting predictions retained a positive correlation with measured activity on the held-out test set (Spearman $\rho$= 0.53 for all variants and $\rho$ = 0.50 for variants with activity $\geqslant$WT; **Figure S3a**). The position-specific rankings were identical to those obtained with all 11 plates at six of the seven positions. Although the ranking at position 59 changed because they had similar predicted values in both cases, all four available amino acids at this position were retained and therefore did not affect the final library composition (**Figure S3b**).

Applying the same per-position library sizes produced the same 144-variant MLEE-enriched library from both models: F/I/K/T59, F/I/L63, F154, Y/L156, G157, A/S159, and A/F/L161. The predicted activity magnitudes differed between the models, but the amino-acid selections determining the library remained unchanged (**Figure S4**). Thus, for MthUPO, four initial screened plates provided sufficient information to reproduce the library-design decision obtained using the complete 11-plate dataset, indicating that the initial experimental workload could potentially have been reduced without changing the selected MLEE-enriched library.

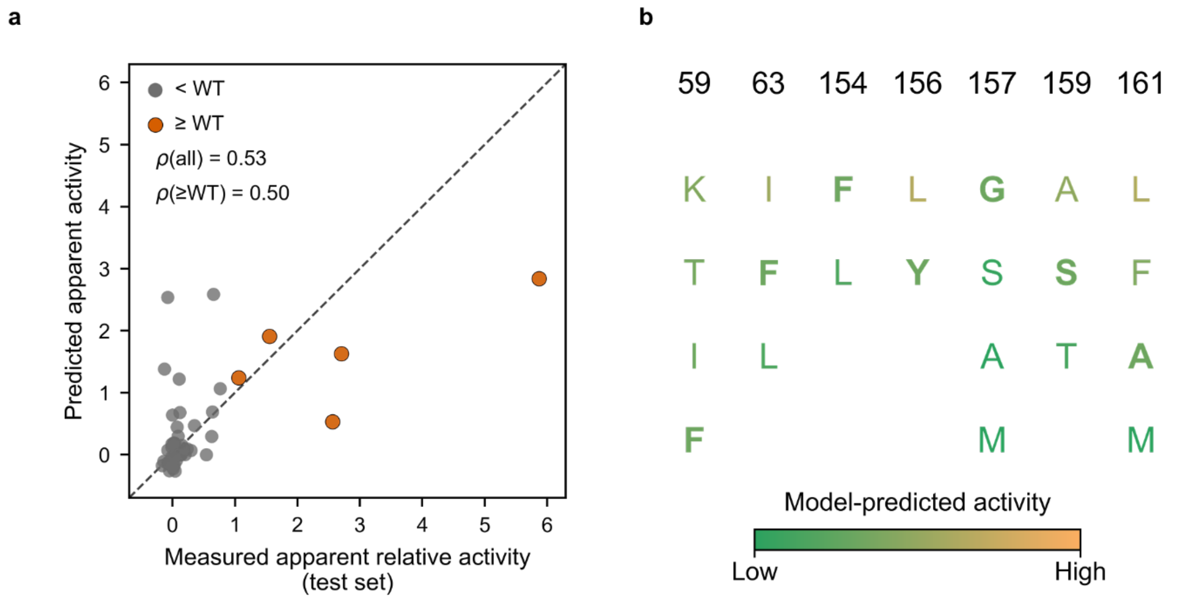


**Figure S3. Model performance and position-specific amino-acid ranking using four initial screening plates**. **a. Predicted versus measured apparent relative activity for the held-out test set**. Gray and orange points indicate variants with activity below and at or above WT, respectively; ρ denotes the Spearman correlation coefficient. **b. Position-specific amino-acid rankings based on model-predicted activity**. Amino acids are ordered from highest to lowest predicted activity at each position, and WT identities are shown in bold.

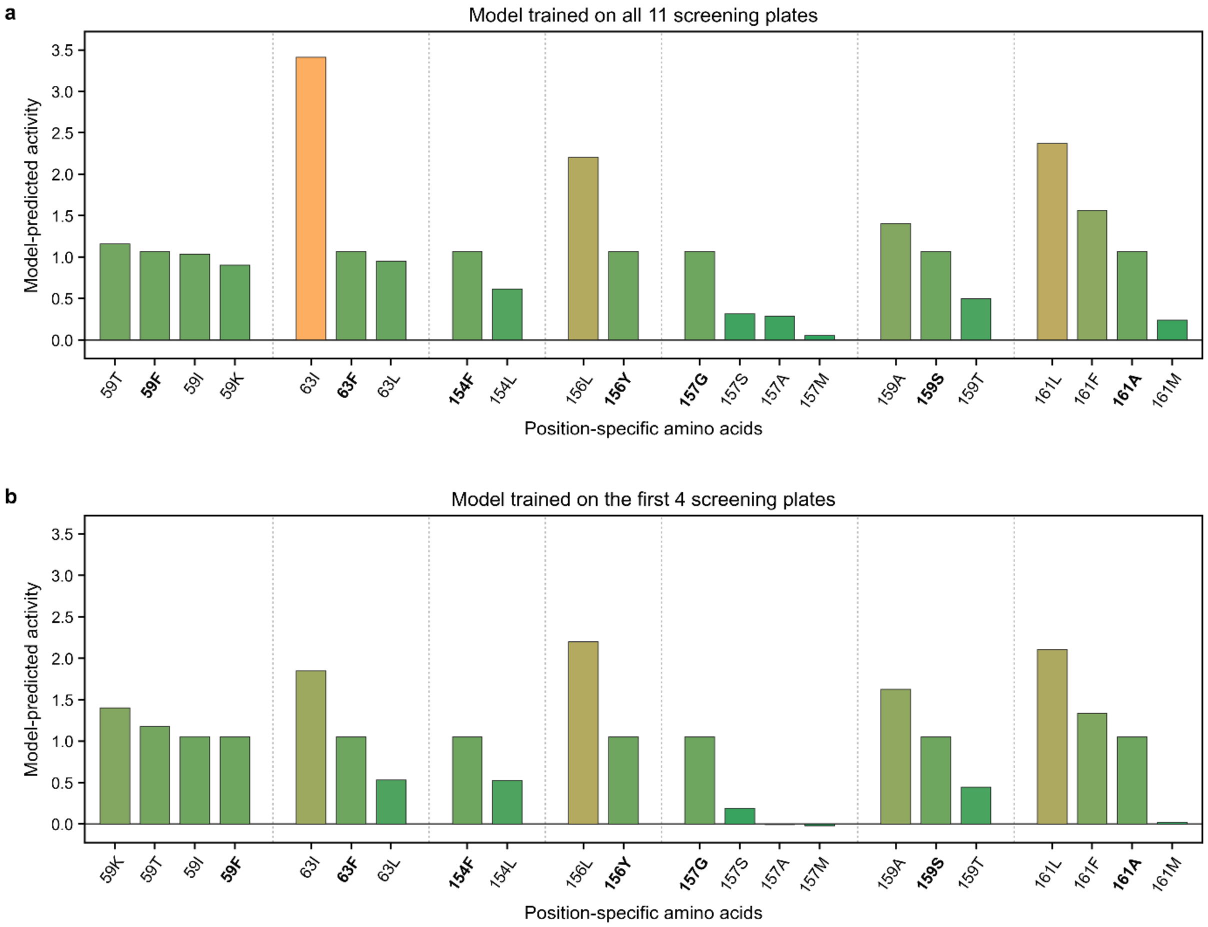


**Figure S4**. **Comparison of position-specific amino-acid predictions from four and 11 initial screening plates**. Model-predicted activities of single variants obtained using **a. all 11 screened plates** and **b. the first four screened plates**. Within each position, amino acids are ordered from highest to lowest predicted activity; WT identities are shown in bold. Both models supported the same 144-variant MLEE-enriched library composition.

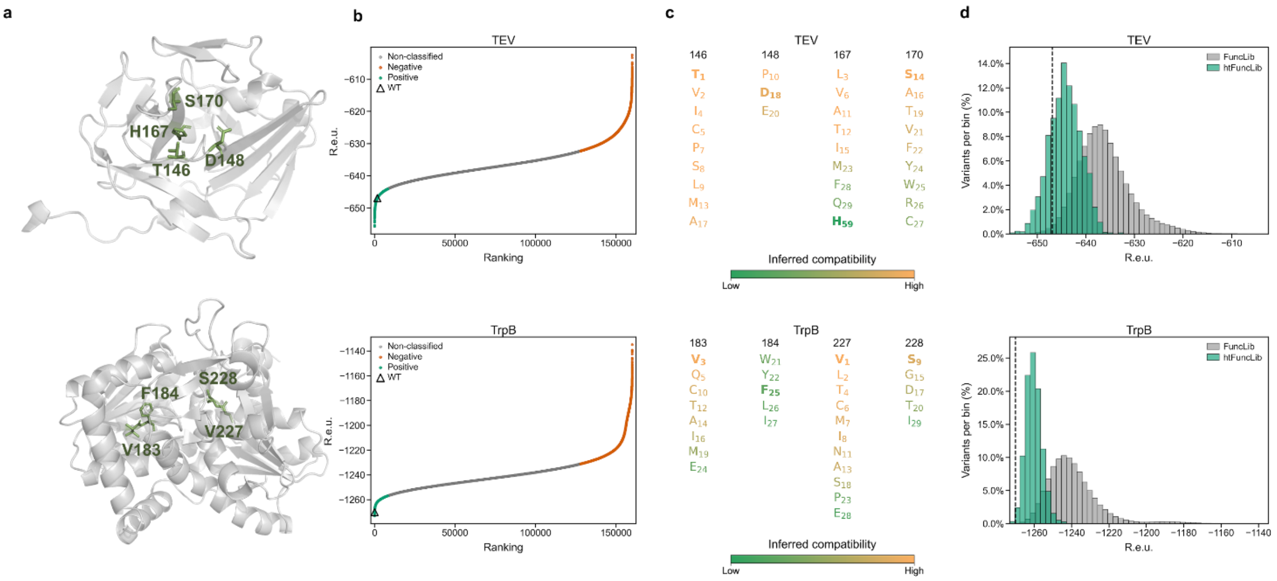


**Figure S5.** The same as **Figure 2**. The corresponding PDB structures for TEV and TrpB were 1LVM, and 8VHH.

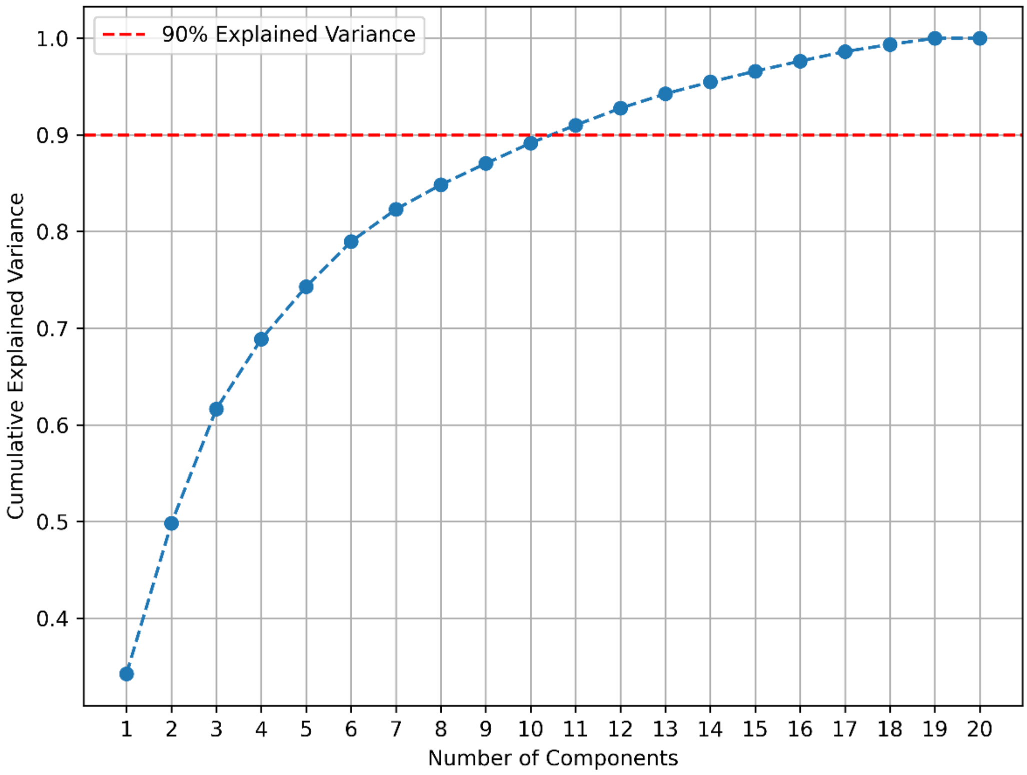


**Figure S6. Cumulative explained Variance of PCA.**

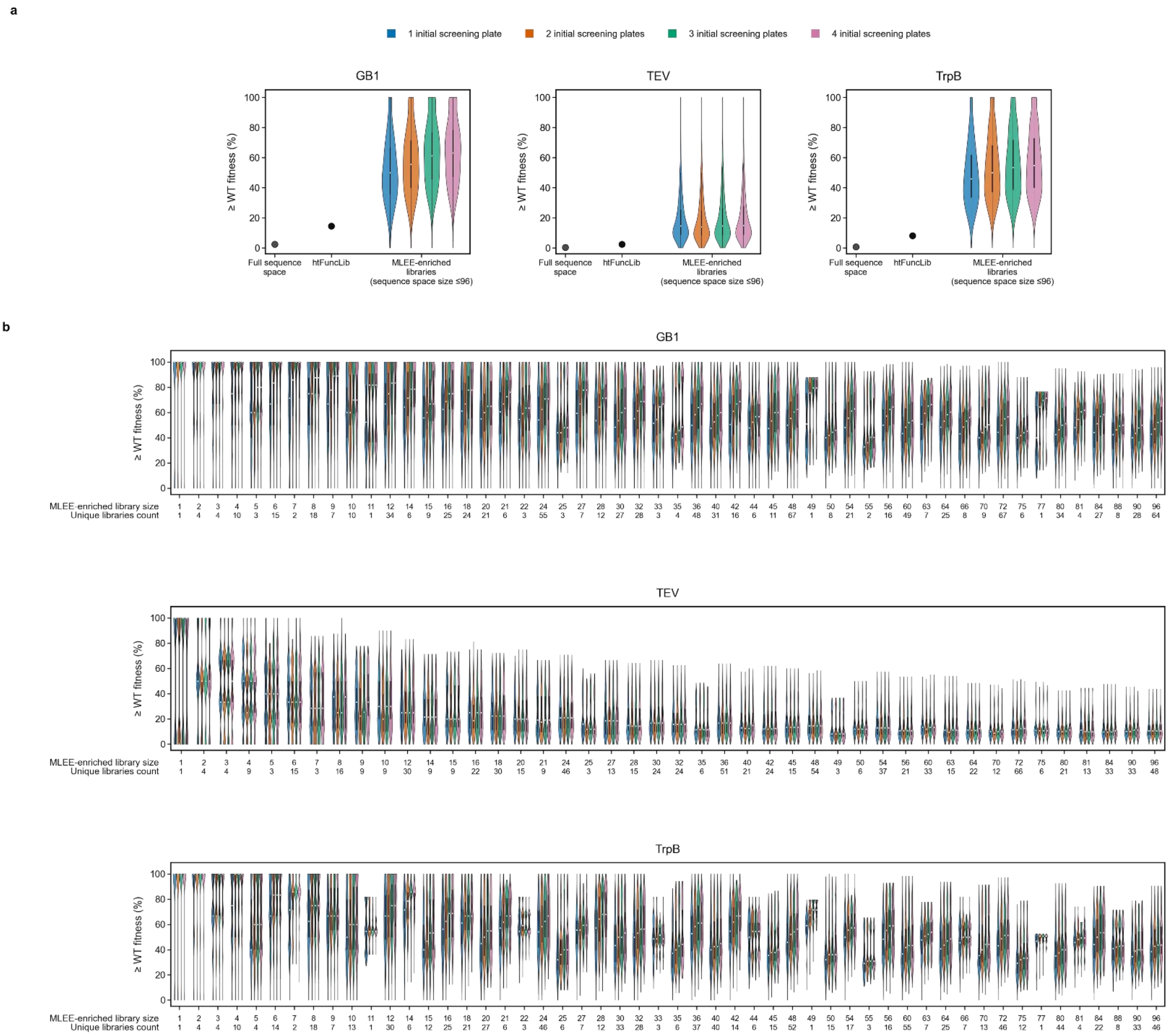


**Figure S7**. **Distribution of percentages of variants at or above WT fitness for all MLEE-enriched libraries with sequence space ≤ 96, in general (a), and in exact library size (b), across all simulations.**

.

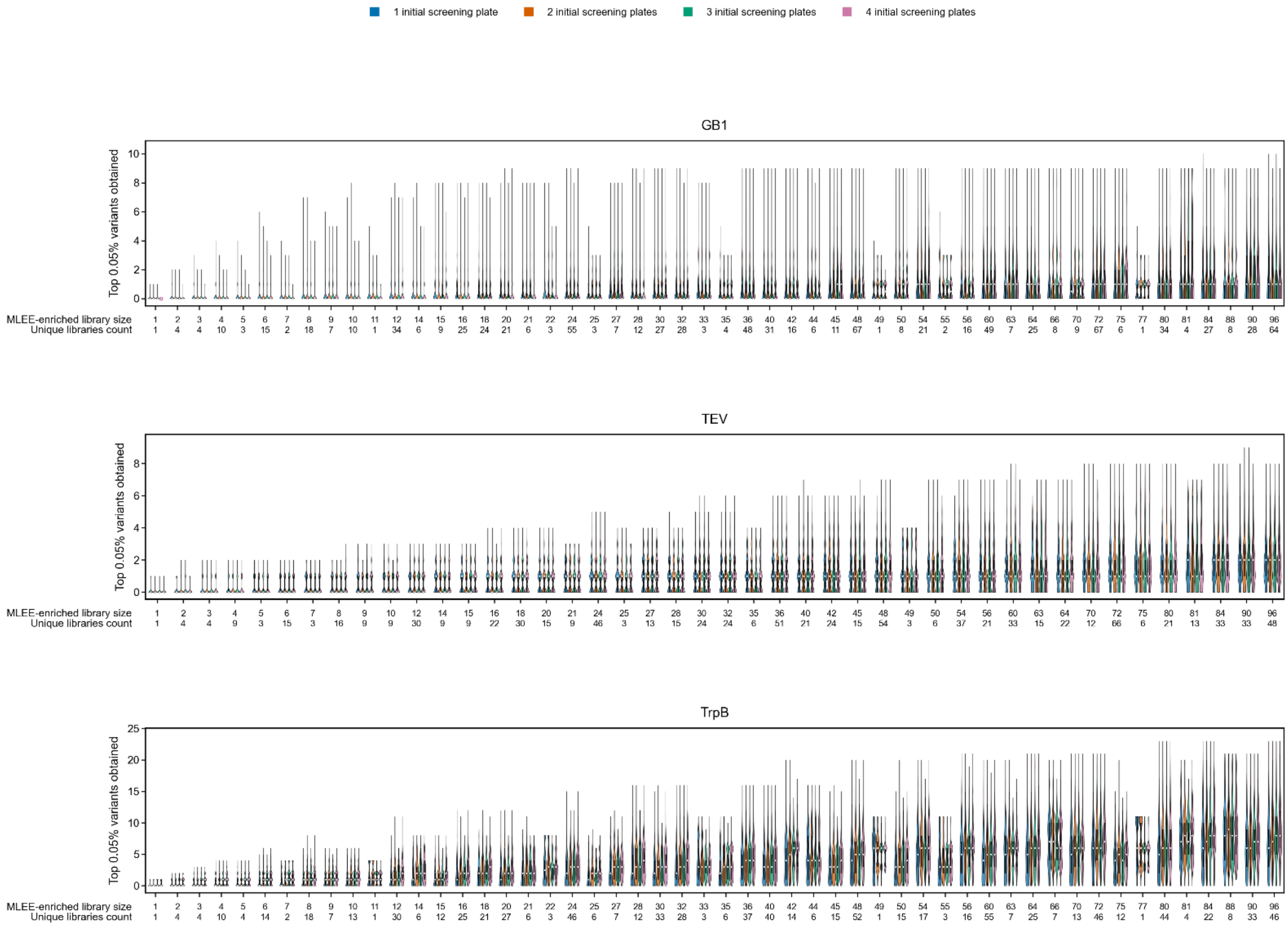


**Figure S8. Distribution of top 0.05% variants obtained in all MLEE-enriched libraries with sequence space ≤ 96.**

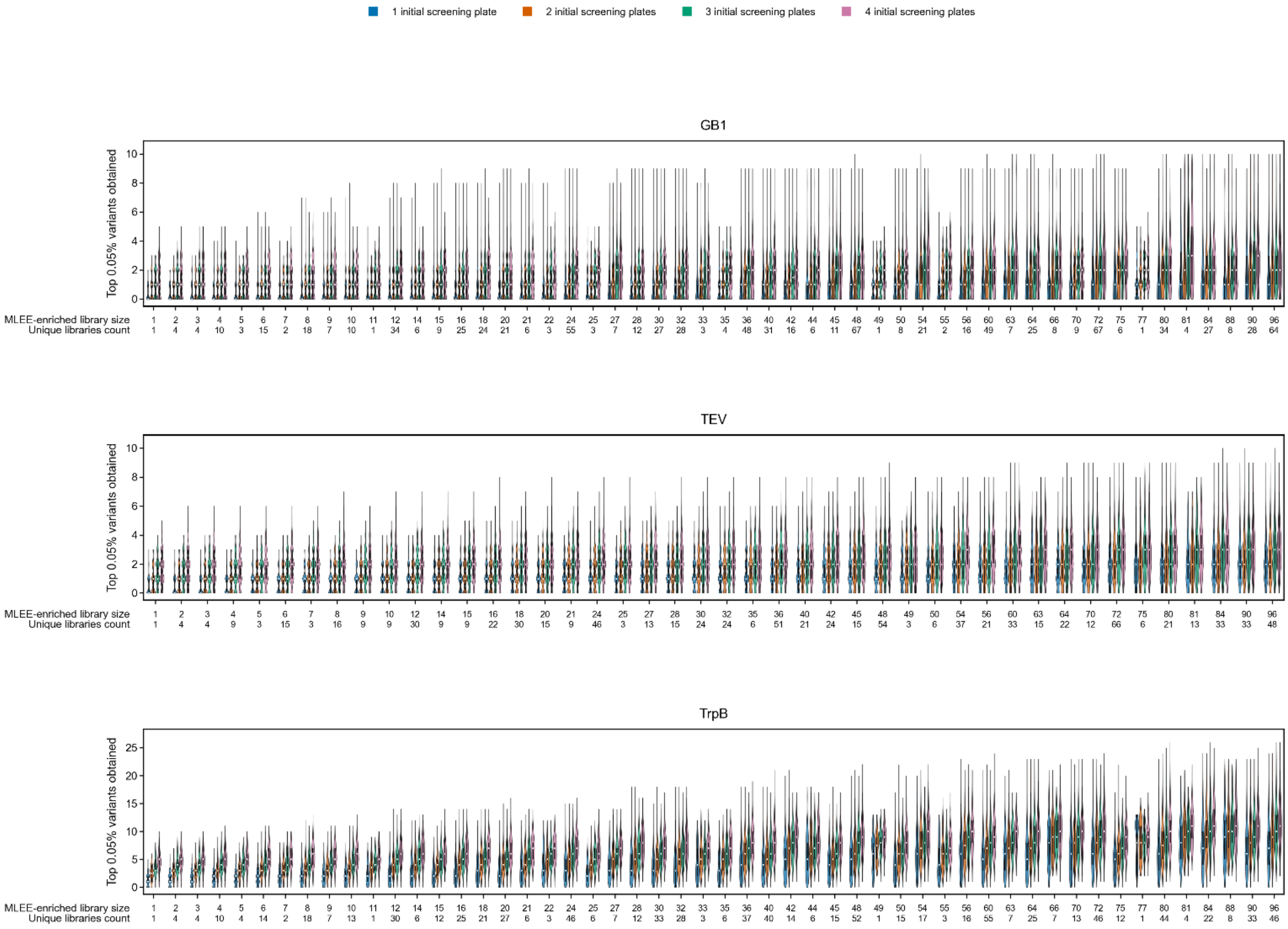


**Figure S9. Distribution of top 0.05% variants obtained when combine initial round screenings and MLEE-enriched libraries with sequence space ≤ 96.**

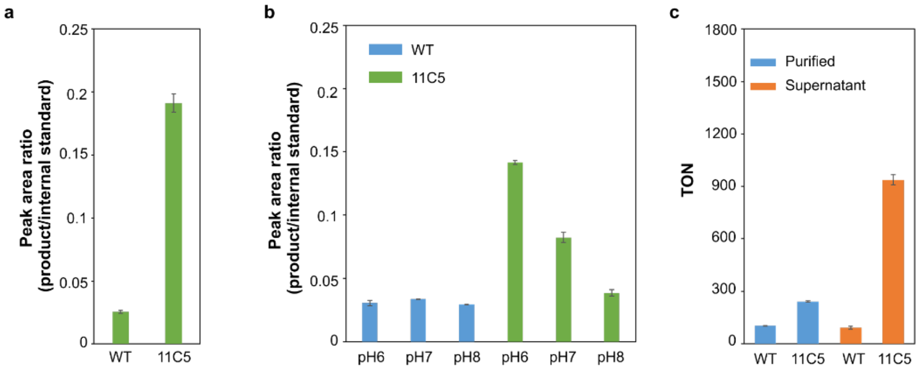


**Figure S10 TON assay optimization. a. Activity of purified enzymes in heated-supernatant.** Respective supernatants with expressed enzymes were first heat-deactivated at 95 °C for 0.5 h, and then were used to dilute the correspondent enzymes to 500 mM before adding 100 μL to the 400 uL reaction mastermix to do the reaction. Negative control with heat-deactivated supernatants were assayed in the same way with no activity. All samples were in duplicate. **b. Activity of purified enzymes in different pH.** 500 uL reaction contains 100 nM respective enzyme under different pH conditions. All sample in duplicate. **c. TON of purified enzymes and supernatants.** For purified enzymes, the enzyme concentrations in reaction mixture were 100 nM for both. For supernatants, the concentrations were 62 nM for WT and 68 nM for 11C5. All samples in triplicate**.**

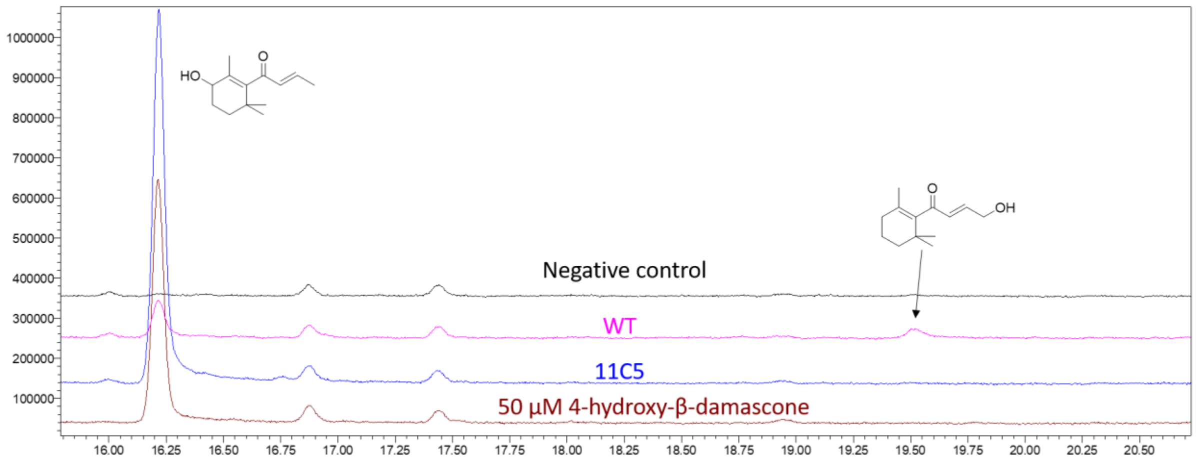


**Figure S11 Regioselectivity of WT and 11C5.** Total ion chromatography of WT and 11C5 from the same experiment of Fig 5a. 10-hydroxy-β-damascone was determined according to ref[3].

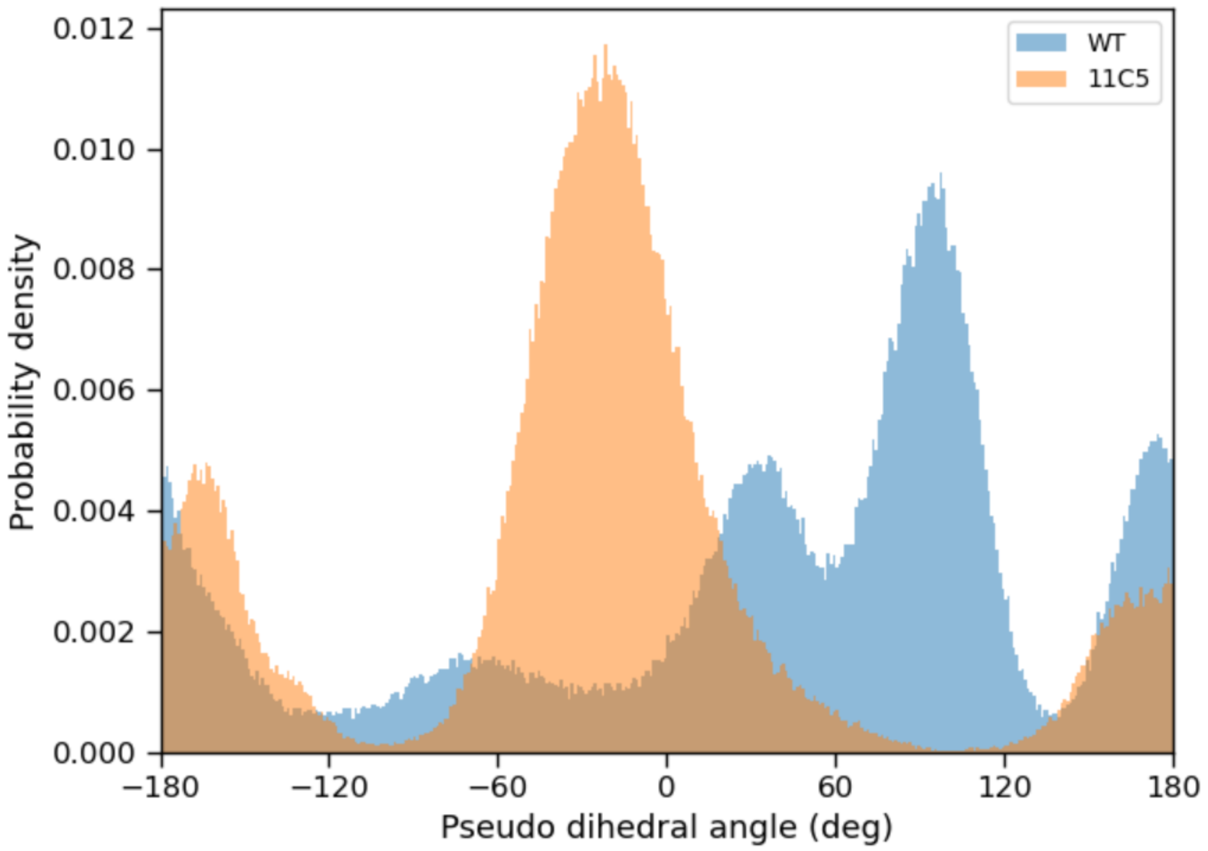


**Figure S12. Probability density distributions of the pseudo-dihedral angle formed by atoms Fe–O–C4–C5 in WT and 11C5.** The shifts in angular sampling reflect distinct substrate orientations relative to the reactive $Fe^{IV}$=O center in the two variants. Notably, 11C5 samples a more well-defined substrate conformation compared with the WT.

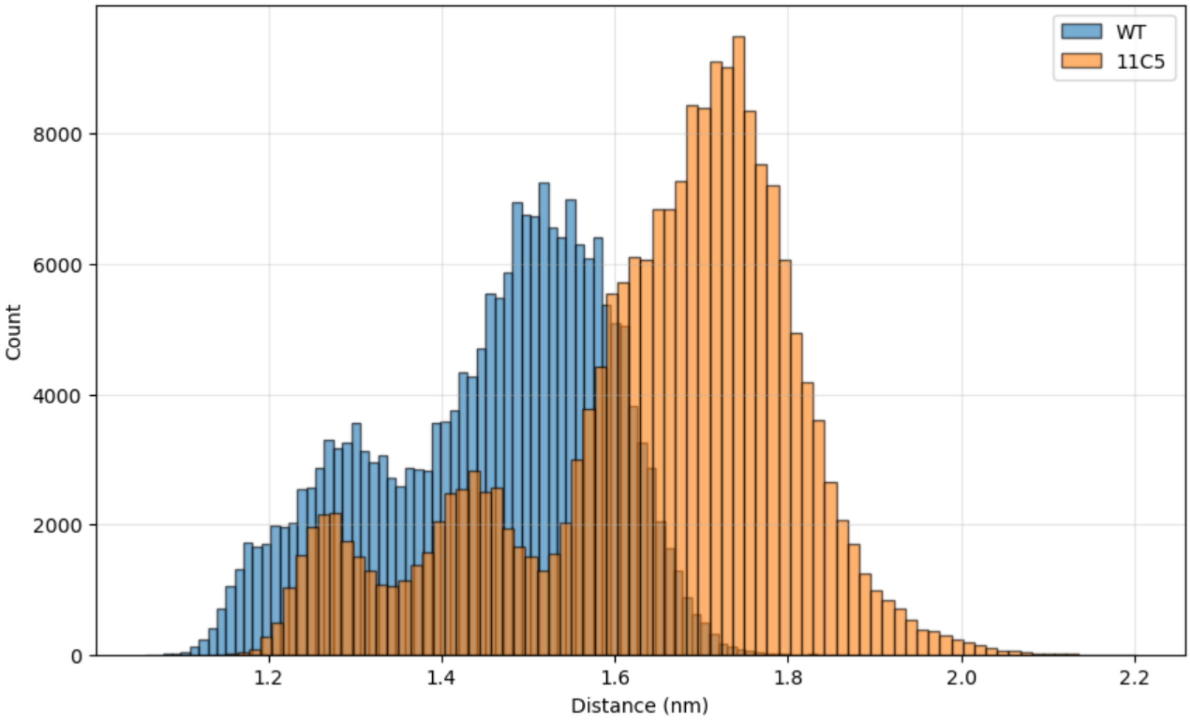


**Figure S13. Distance distributions between helix A and helix B for WT and 11C5**. Distances were obtained by computing the center-of-mass (COM) separation between the CA atoms of residues 63–68 (helix A) and residues 149–165 (helix B), capturing the relative opening of the substrate-access channel in 11C5 compared with the WT.

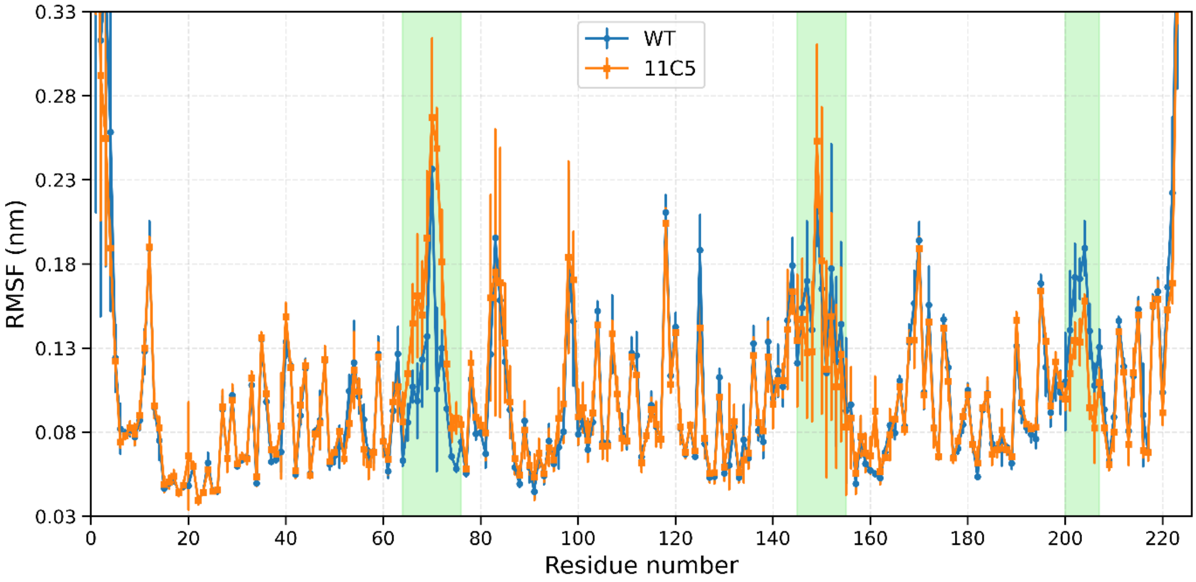


**Figure S14 Residue-wise RMSF comparison between WT and 11C5 derived from five independent MD replicas.** The green highlighted regions correspond to secondary-structure elements displaying the strongest shifts in flexibility—particularly helix A(residues 64 to 76), helix B(residues 145 to 155), and helix C(residues 200 to 207)—revealing distinct dynamic signatures between the two variants.

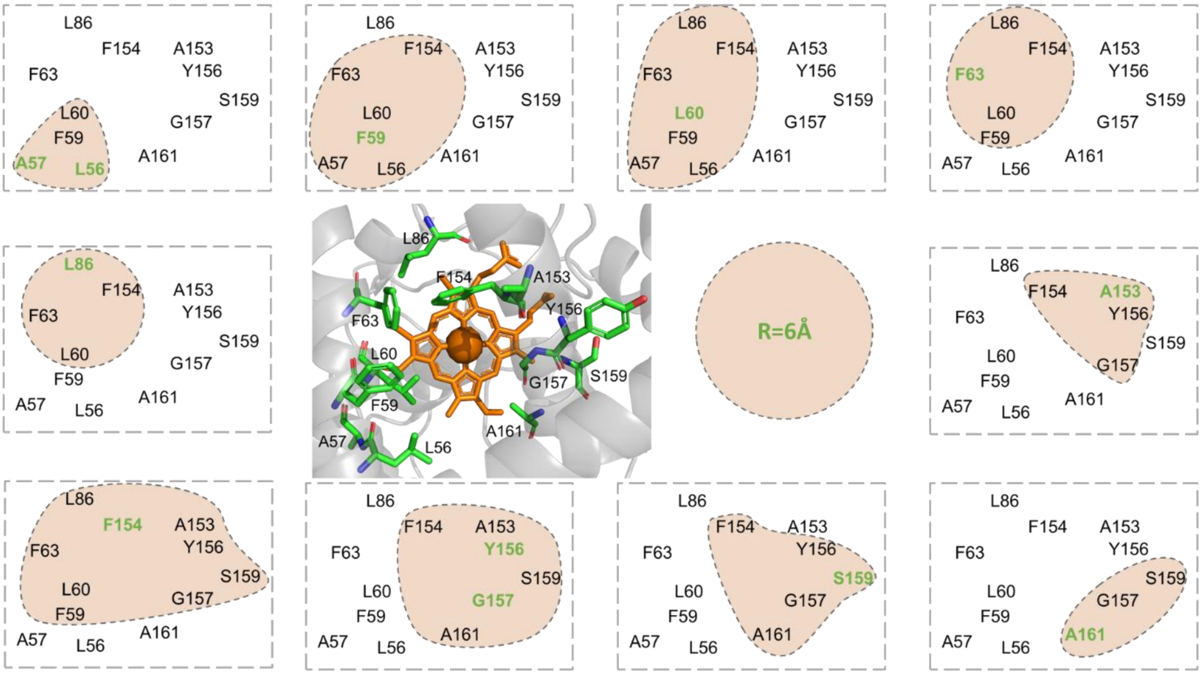


**Figure S15. Adjacent neighbourhoods for 2$^{nd}$ FuncLib calculation.** 10 adjacent neighbourhoods were constructed according to a

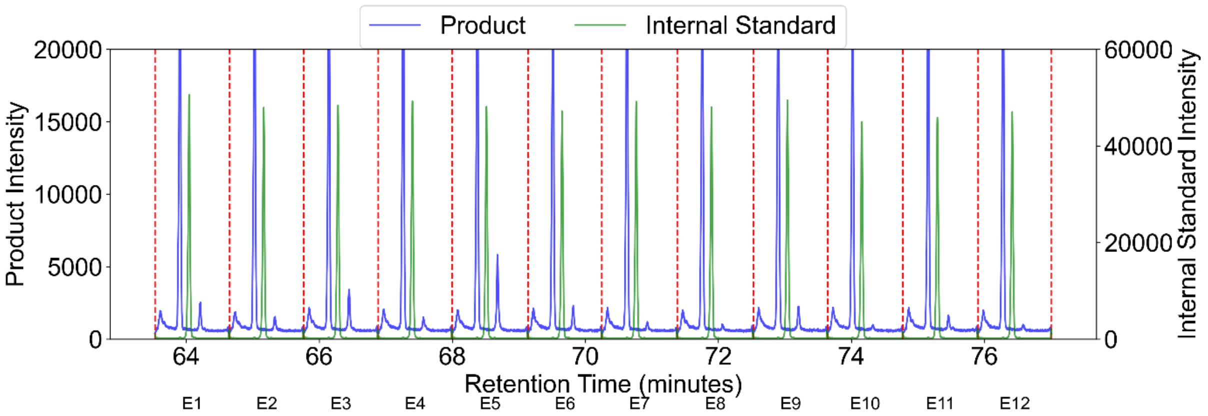


**Figure S16. A snippet of the MISER GC-MS chromatograph.** Product peak is m/z 193.00, internal standard peak is m/z 189.00. E6 is from WT.

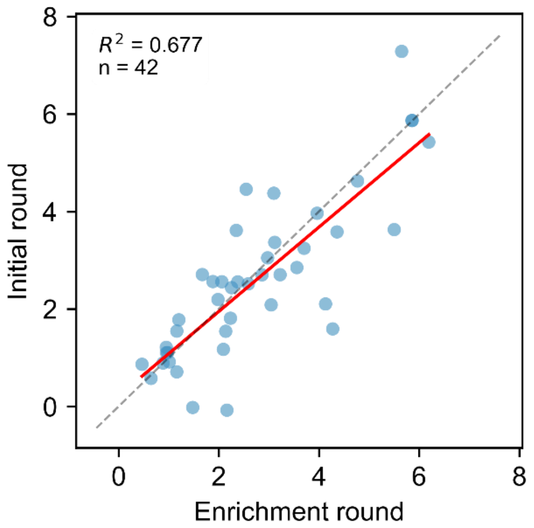


**Figure S17. Calibration between initial round screening and activity enriched library screening.** y=0.22 + 0.87x, $R^2$=0.68

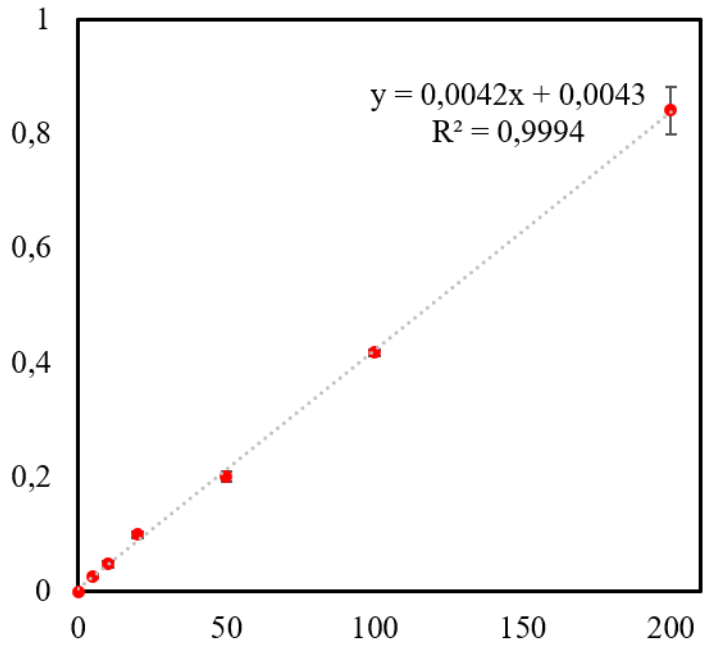


**Figure S18 Calibration curve of 4-hydroxy-β-damascone**

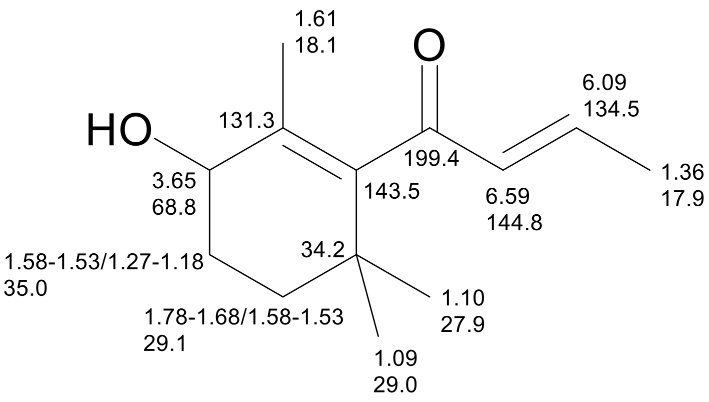

1.61
18.1
O
6.09
134.5
HO
131.3
199.4
1.36
17.9
3.65
68.8
143.5
6.59
144.8
1.58-1.53/1.27-1.18
35.0
34.2
1.10
27.9
1.78-1.68/1.58-1.53
29.1
1.09
29.0


### $^1$H-NMR

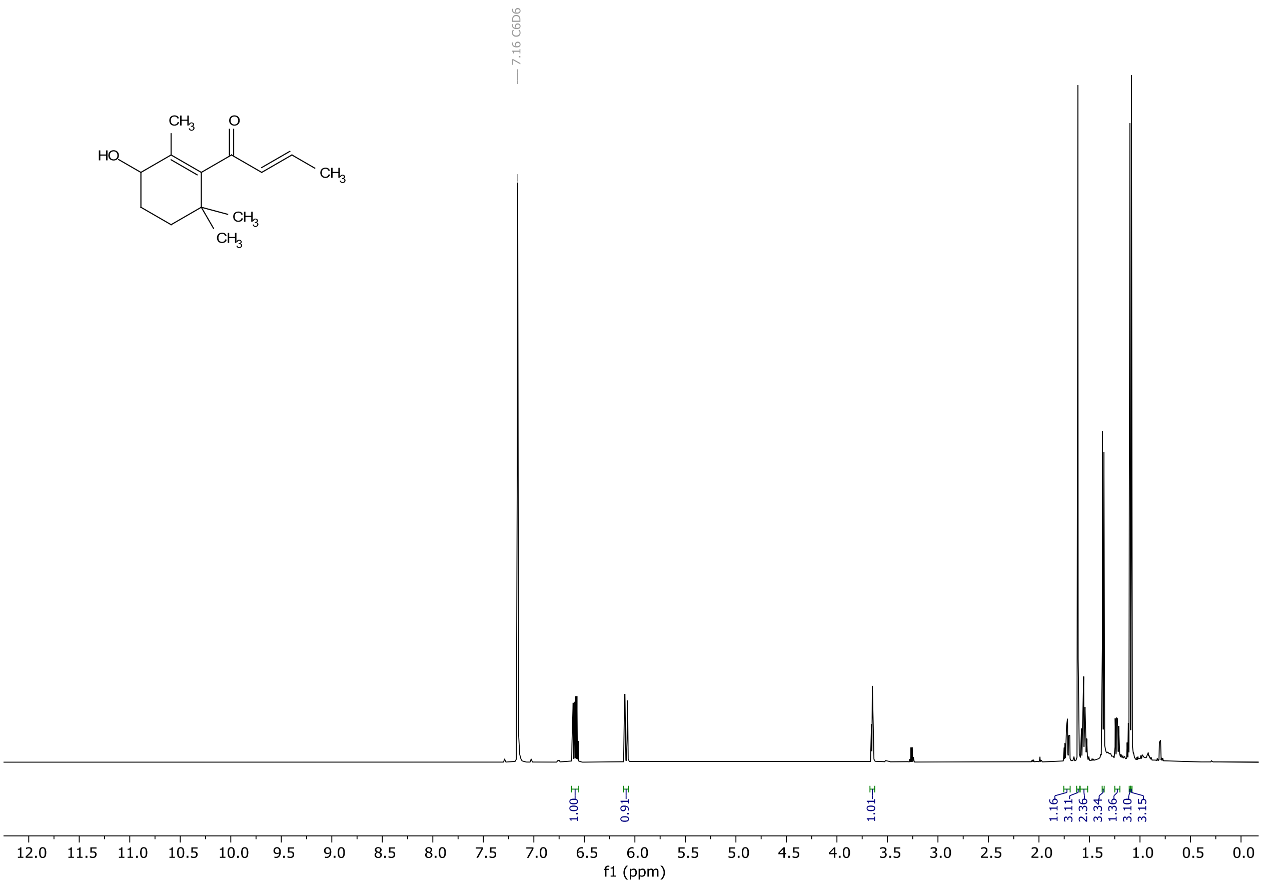

7.16 C6D6
$CH_3$
O
HO
$CH_3$
$CH_3$
$CH_3$
1.00
0.91
1.01
1.16
3.11
2.36
3.34
1.36
3.10
3.15
12.0 11.5 11.0 10.5 10.0 9.5 9.0 8.5 8.0 7.5 7.0 6.5 6.0 5.5 5.0 4.5 4.0 3.5 3.0 2.5 2.0 1.5 1.0 0.5 0.0
f1 (ppm)

### $^{13}C$-NMR

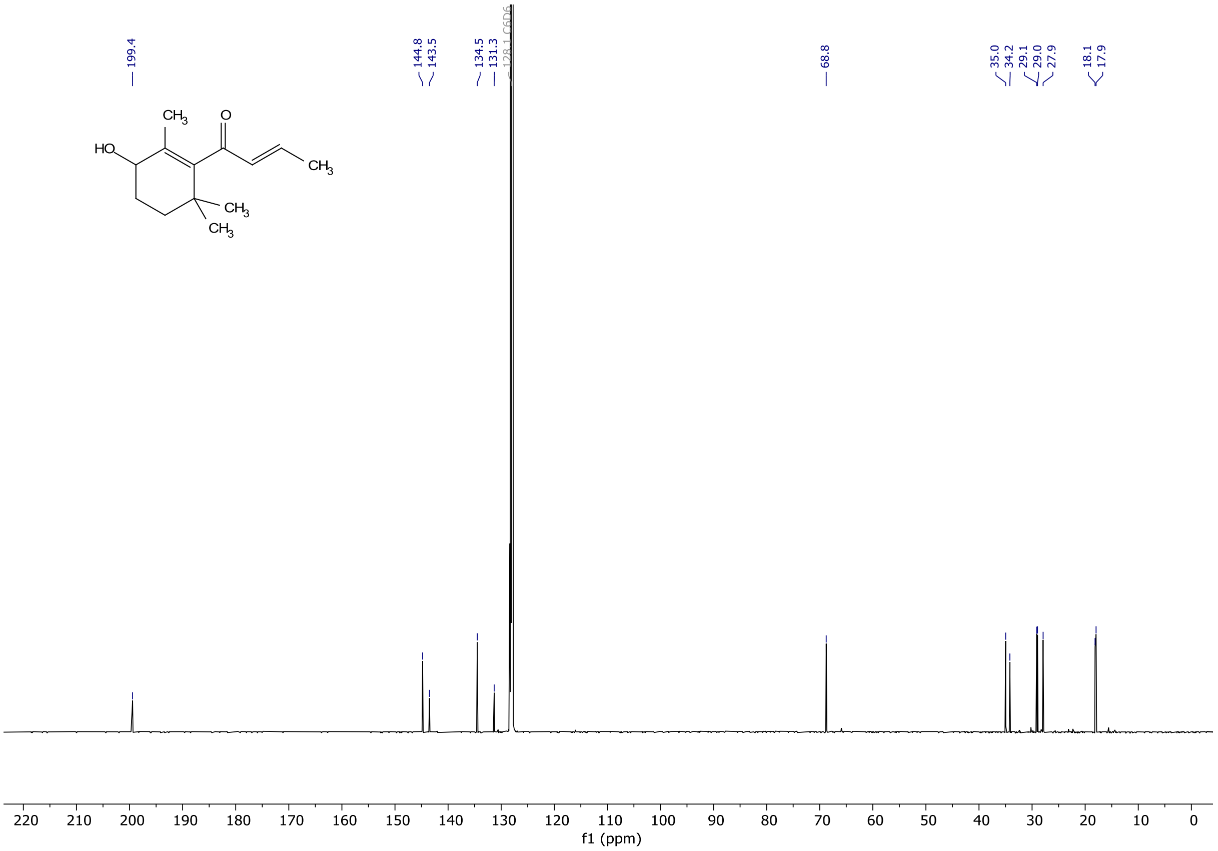


### $^{1}H$-$^{1}H$ COSY

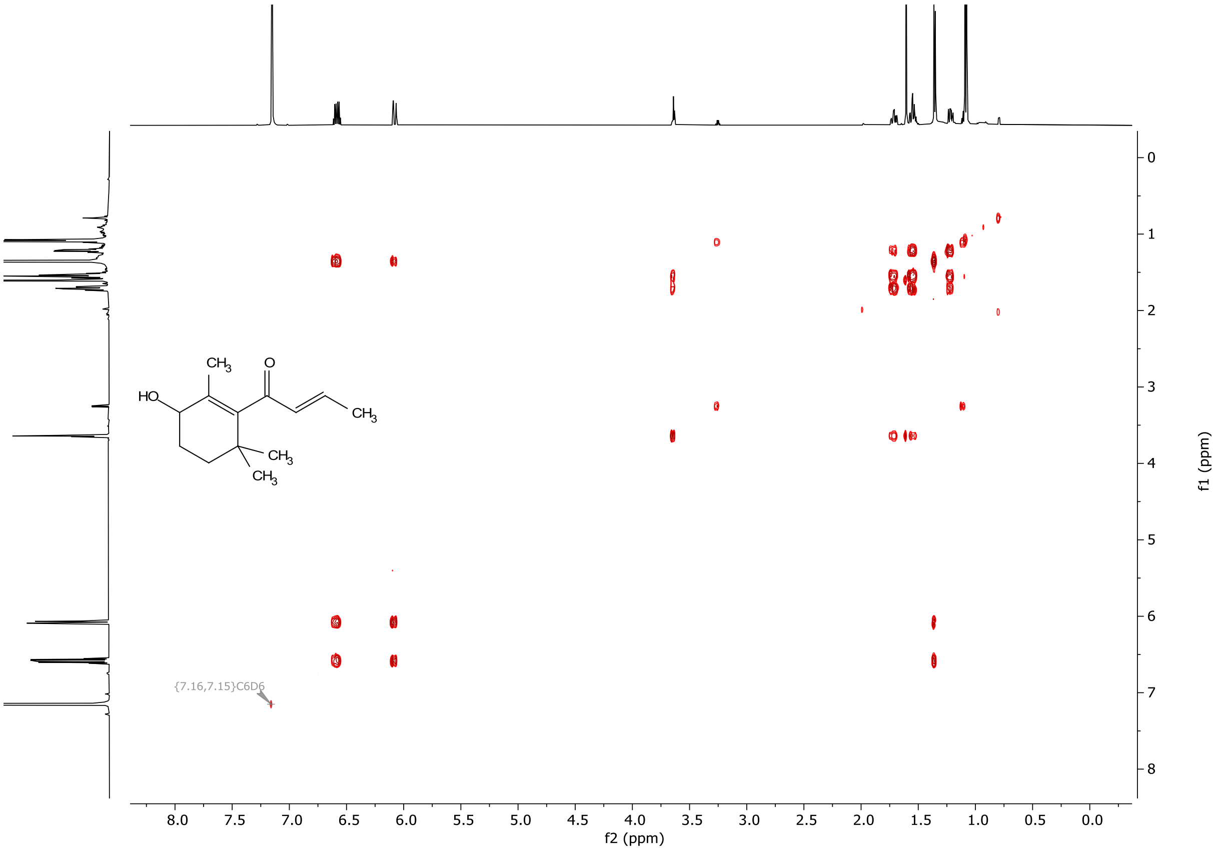

$^{1}H$-$^{13}C$ HSQC

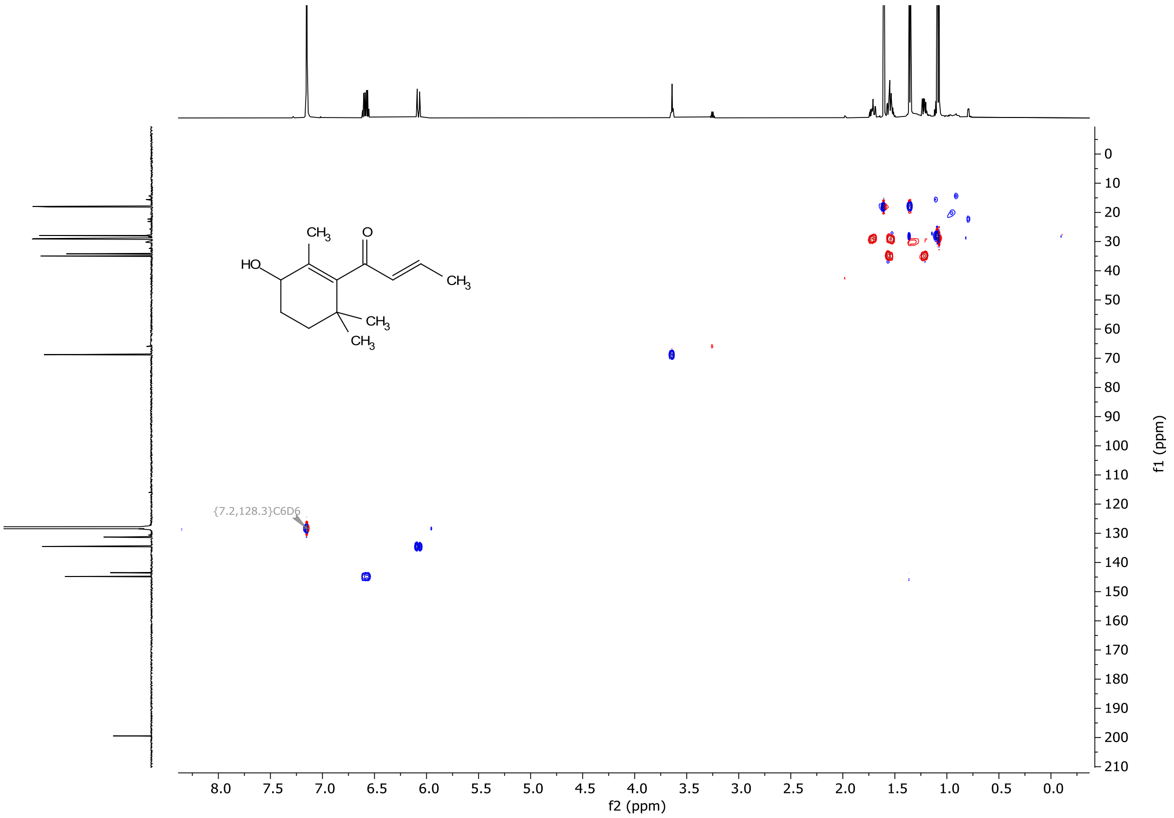


$^{1}H$-$^{13}C$ HMBC

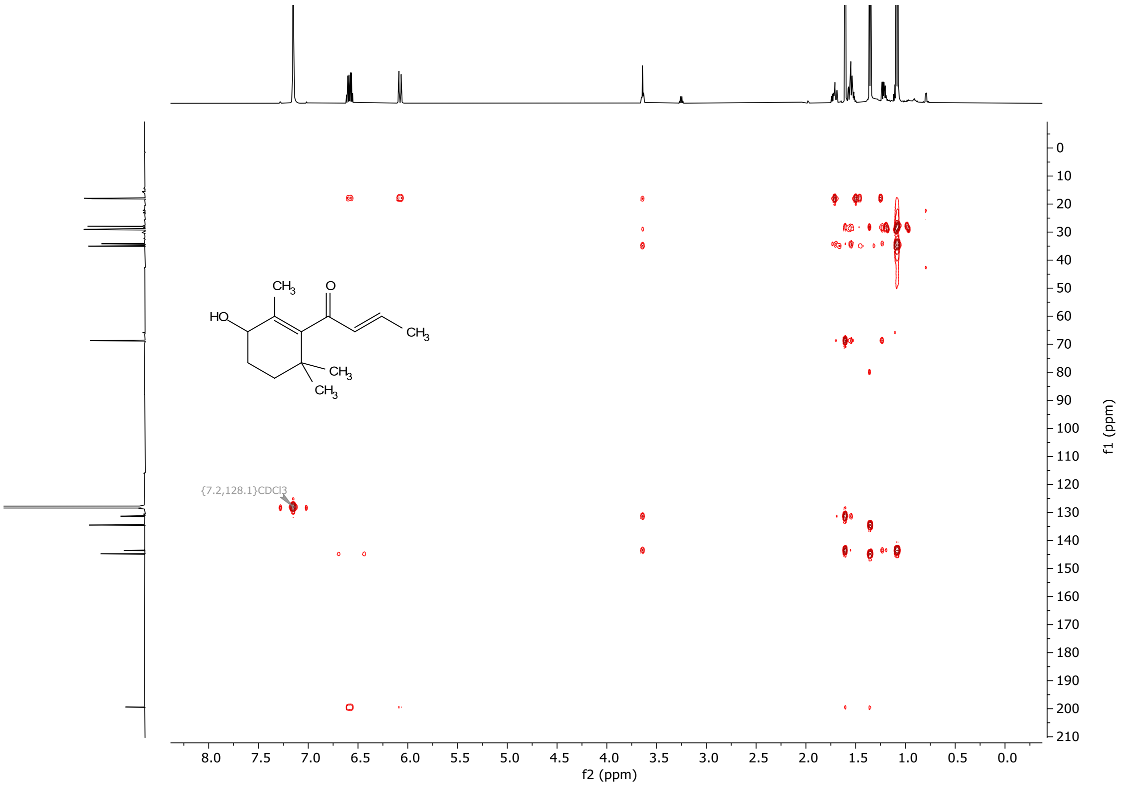


**Figure S19 NMR characterization of 4-hydroxy-β-damascone**

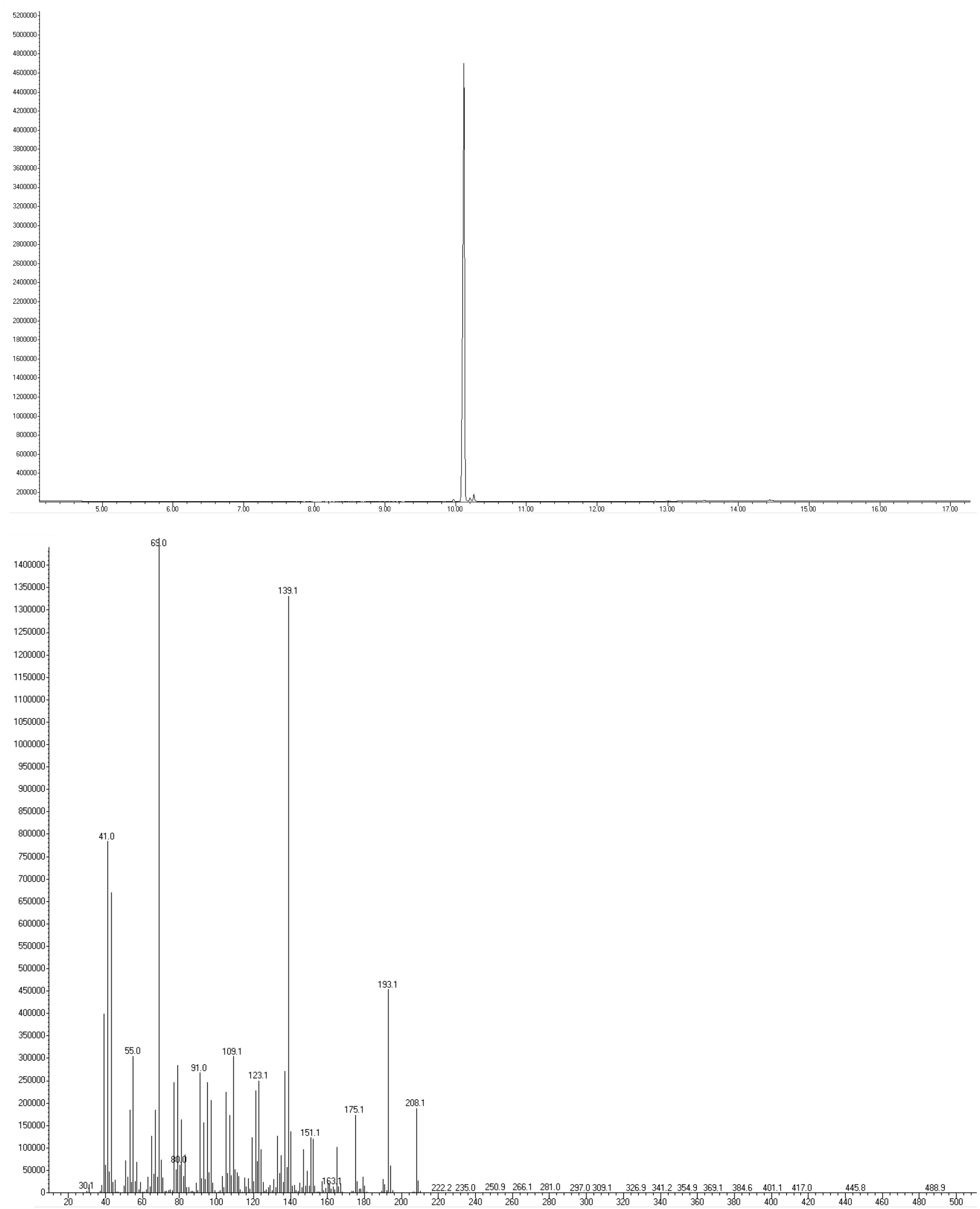


**Figure S20 GC-MS characterization of 4-hydroxy-β-damascone**

**Table S1 Benchmark datasets**

| Protein | Function | Target positions | Full sequence space size | Variants with fitness value | WT fitness | ≥ WT (%) | Count of top 0.05% | Top 0.05% fitness threshold | Max fitness |
|---|---|---|---|---|---|---|---|---|---|
| GB1[4] | Binding affinity to IgG-Fc | V39/D40/G41/V54 | 160000 | 149361 | 0.114 | 2.44 | 75 | 0.597 | 1 |
| TEV[5, 6]* | Protease activity to ENLYFQ↓S | T146/D148/H167/S170 | 160000 | 151827 | 0.295 | 0.21 | 76 | 0.345 | 0.673 |
| TrpB[7] | Tryptophan synthesis | V183/F184/V227/S228 | 160000 | 159129 | 0.408 | 0.69 | 80 | 0.688 | 1 |

* Whenever fitness value is considered, only consider those in original dataset with sequence reads ≥ 5.

**Table S2 htFuncLib libraries of benchmark proteins**

| Protein | Position 1 | Position 2 | Position 3 | Position 4 | Sequence space size | Variants with fitness value | Counts ≥WT (Ratio%) | Max Fitness | Median Fitness | Counts in top 0.05% |
|---|---|---|---|---|---|---|---|---|---|---|
| GB1[4] | VLEQINT SDMAC | DYNF | GSATPC | VAGSMIL C | 2304 | 2231 | 322 (14.4%) | 0.863 | 0.002 | 10 |
| TEV[5, 6]* | TVICPSL MA | DPE | HLVATIM FQ | SATVFY WRC | 2187 | 2139 | 51 (2.38%) | 0.404 | 0.001 | 10 |
| TrpB[7] | VQCTAIM E | FWYLI | VLTCMIN ASPE | SGDTI | 2200 | 2195 | 177 (8.06%) | 0.833 | 0.023 | 30 |

* Whenever fitness value is considered, only consider those in original dataset with sequence reads ≥ 5.

**Table S3 AAindex_PC11 descriptors**

| AA | PC1 | PC2 | PC3 | PC4 | PC5 | PC6 | PC7 | PC8 | PC9 | PC10 | PC11 |
|---|---|---|---|---|---|---|---|---|---|---|---|
| **A** | -0,12034 | -5,42584 | 17,01675 | 2,204228 | 2,819656 | 3,031397 | -3,77326 | 0,45196 | 3,968869 | -1,78279 | -0,80297 |
| **C** | 8,227071 | -8,15197 | -6,01551 | -14,3638 | 12,44592 | -7,31302 | -3,36786 | 5,209335 | -1,9149 | 4,099915 | -2,0778 |
| **D** | -18,5022 | 3,041214 | -0,07492 | -1,68545 | 5,738213 | 5,063807 | 9,413391 | 2,1688 | -1,97 | -1,62867 | -1,51653 |
| **E** | -12,1659 | 11,91135 | 10,55553 | 4,395216 | 7,926747 | 4,113799 | 4,766318 | 3,686365 | -2,37204 | -1,41992 | 0,496874 |
| **F** | 19,68816 | 0,372806 | -3,94566 | 1,698549 | -1,95706 | 4,111803 | 1,089936 | -2,30192 | -1,2772 | -1,4164 | -3,71816 |
| **G** | -16,8312 | -21,6292 | 3,123578 | -7,33567 | -9,42616 | 8,832825 | -4,84776 | 3,544309 | -3,04697 | -2,23371 | 1,946356 |
| **H** | -0,33574 | 8,323947 | -4,66385 | -4,97241 | 1,084526 | 0,702253 | -3,41945 | -9,5475 | -2,8214 | -3,56804 | -0,82481 |
| **I** | 21,01489 | -5,85756 | 2,403985 | 4,249276 | -2,48449 | -2,77021 | 3,875807 | 0,35742 | -3,94945 | 0,406186 | 3,829768 |
| **K** | -11,8128 | 13,54179 | 5,459675 | 1,929035 | -5,95619 | -2,46158 | -4,36055 | 0,392793 | -0,11167 | 8,994955 | 4,238633 |
| **L** | 17,91927 | -3,0869 | 11,30729 | 6,859729 | -2,10782 | 2,222848 | -0,39899 | -0,92874 | 0,441807 | 5,341424 | -7,46657 |
| **M** | 16,16844 | 4,852779 | 3,45758 | -2,00032 | 6,55925 | 3,373442 | -6,62034 | -4,00504 | -1,03055 | -3,31099 | 3,208925 |
| **N** | -15,4278 | 0,095906 | -3,61245 | -6,31184 | -1,59956 | 1,967577 | 3,610538 | -5,55857 | -1,10098 | 4,612496 | -2,20575 |
| **P** | -16,5046 | -11,8824 | -16,929 | 19,22077 | 6,085233 | -1,03288 | -4,80778 | -0,1061 | -0,98251 | 0,481937 | -0,30673 |
| **Q** | -7,97785 | 9,140132 | 1,733519 | -0,06425 | 1,715818 | -2,13458 | -1,49976 | -0,57799 | 0,631208 | 0,14547 | 5,224574 |
| **R** | -8,32446 | 15,80111 | -1,26794 | 0,054628 | -8,12278 | -8,34681 | -4,80263 | 5,875141 | -1,43613 | -5,79495 | -5,71715 |
| **S** | -12,7683 | -7,99236 | 1,63439 | -3,08549 | -1,30996 | -3,99018 | 2,017706 | -3,40154 | 6,01126 | 1,042786 | -1,75995 |
| **T** | -4,83891 | -5,97115 | -0,06309 | -0,56231 | -1,27589 | -7,64977 | 4,13978 | -2,47983 | 7,699965 | -3,7482 | 1,488047 |
| **V** | 16,19321 | -8,56657 | 5,880621 | 2,613782 | -1,86403 | -7,6502 | 4,306114 | 1,470467 | -2,32489 | -1,69706 | 3,350785 |
| **W** | 17,82514 | 8,438268 | -13,0671 | -1,64422 | -0,60607 | 9,140289 | -0,34633 | 5,481274 | 8,42173 | 0,575093 | 1,64695 |
| **Y** | 8,573888 | 3,044662 | -12,9334 | -1,19948 | -7,66535 | 0,78919 | 5,025134 | 0,269374 | -2,83614 | 0,900454 | 0,965502 |

**Table S4** Sequence space for htFuncLib calculation

| Pos. | WT | 1st FuncLib | Previous work[8-10] |
|---|---|---|---|
| 56 | L | V | I |
| 57 | A | C | I |
| 59 | F | AHIKLRSTVY | Q |
| 60 | L | M | QFI |
| 63 | F | AEHILQVY | |
| 86 | L | M | IV |
| 153 | A | DEHIKLMNQRSTV | |
| 154 | F | LM | IV |
| 156 | Y | FLRV | I |
| 157 | G | AFLMST | |
| 159 | S | ACTV | NGY |
| 161 | A | FLM | VI |
| **Seq. space** | | **2743372800** | |

**Table S5** Point mutations ranked by EpiNNet

| Rank | position | AA | Neg. poss | Pos. poss | Rank | position | AA | Neg. poss | Pos. poss |
|---|---|---|---|---|---|---|---|---|---|
| **1** | **161** | **F** | **0.000** | **1.000** | 42 | 153 | R | 1.000 | 0.000 |
| **2** | **157** | **A** | **0.004** | **0.996** | 43 | 63 | F | 1.000 | 0.000 |
| **3** | **157** | **M** | **0.011** | **0.989** | 44 | 153 | V | 1.000 | 0.000 |
| **4** | **154** | **L** | **0.068** | **0.932** | 45 | 159 | C | 1.000 | 0.000 |
| **5** | **159** | **A** | **0.078** | **0.922** | 46 | 159 | V | 1.000 | 0.000 |
| **6** | **161** | **M** | **0.119** | **0.881** | 47 | 59 | H | 1.000 | 0.000 |
| **7** | **159** | **T** | **0.207** | **0.793** | 48 | 59 | L | 1.000 | 0.000 |
| **8** | **161** | **L** | **0.333** | **0.667** | 49 | 156 | V | 1.000 | 0.000 |
| **9** | **59** | **K** | **0.338** | **0.662** | 50 | 156 | R | 1.000 | 0.000 |
| **10** | **63** | **L** | **0.604** | **0.396** | 51 | 59 | R | 1.000 | 0.000 |
| **11** | **156** | **L** | **0.635** | **0.365** | 52 | 63 | E | 1.000 | 0.000 |
| **12** | **157** | **S** | **0.937** | **0.063** | 53 | 153 | S | 1.000 | 0.000 |
| **13** | **59** | **T** | **0.949** | **0.051** | 54 | 154 | I | 1.000 | 0.000 |
| **14** | **59** | **I** | **0.953** | **0.047** | 55 | 153 | N | 1.000 | 0.000 |
| 15 | 86 | L | 0.962 | 0.038 | 56 | 153 | H | 1.000 | 0.000 |
| 16 | 154 | M | 0.967 | 0.033 | 57 | 156 | I | 1.000 | 0.000 |
| 17 | 60 | L | 0.967 | 0.033 | 58 | 63 | V | 1.000 | 0.000 |
| 18 | 157 | L | 0.969 | 0.031 | 59 | 57 | I | 1.000 | 0.000 |
| 19 | 159 | S | 0.982 | 0.018 | 60 | 56 | V | 1.000 | 0.000 |
| 20 | 161 | A | 0.984 | 0.016 | 61 | 157 | G | 1.000 | 0.000 |
| 21 | 56 | L | 0.990 | 0.010 | 62 | 59 | V | 1.000 | 0.000 |
| 22 | 153 | E | 0.990 | 0.010 | 63 | 57 | C | 1.000 | 0.000 |
| 23 | 153 | K | 0.991 | 0.009 | 64 | 153 | D | 1.000 | 0.000 |
| 24 | 153 | A | 0.994 | 0.006 | 65 | 86 | M | 1.000 | 0.000 |
| 25 | 153 | L | 0.994 | 0.006 | 66 | 63 | H | 1.000 | 0.000 |
| 26 | 59 | Y | 0.997 | 0.003 | 67 | 63 | A | 1.000 | 0.000 |
| 27 | 63 | I | 0.997 | 0.003 | 68 | 59 | A | 1.000 | 0.000 |
| 28 | 60 | M | 0.997 | 0.003 | 69 | 59 | S | 1.000 | 0.000 |
| 29 | 63 | Q | 0.998 | 0.002 | 70 | 159 | Y | 1.000 | 0.000 |
| 30 | 63 | Y | 0.998 | 0.002 | 71 | 59 | Q | 1.000 | 0.000 |
| 31 | 59 | F | 0.998 | 0.002 | 72 | 60 | F | 1.000 | 0.000 |
| 32 | 153 | T | 0.999 | 0.001 | 73 | 159 | G | 1.000 | 0.000 |
| 33 | 156 | Y | 0.999 | 0.001 | 74 | 86 | I | 1.000 | 0.000 |
| 34 | 153 | I | 0.999 | 0.001 | 75 | 159 | N | 1.000 | 0.000 |
| 35 | 157 | T | 0.999 | 0.001 | 76 | 60 | Q | 1.000 | 0.000 |
| 36 | 153 | Q | 0.999 | 0.001 | 77 | 56 | I | 1.000 | 0.000 |
| 37 | 153 | M | 0.999 | 0.001 | 78 | 154 | V | 1.000 | 0.000 |
| 38 | 154 | F | 0.999 | 0.001 | 79 | 86 | V | 1.000 | 0.000 |
| 39 | 57 | A | 0.999 | 0.001 | 80 | 60 | I | 1.000 | 0.000 |
| 40 | 156 | F | 1.000 | 0.000 | 81 | 161 | I | 1.000 | 0.000 |
| 41 | 157 | F | 1.000 | 0.000 | 82 | 161 | V | 1.000 | 0.000 |

**Table S6** Sequence space of initial round library

| Pos. | WT | Mutations |
|---|---|---|
| 59 | F | KTI |
| 63 | F | LI |
| 154 | F | L |
| 156 | Y | L |
| 157 | G | AMS |
| 159 | S | AT |
| 161 | A | FLM |
| **Seq. space** | | **2304** |

**Table S7** Ambiguous codons at each mutated position for the initial round.

| Protein Position* | DNA_position* | Encoded amino acids | Ambiguous codons |
|---|---|---|---|
| 77 | 229 | ['F', 'I', 'K', 'T'] | WTT, AMA |
| 81 | 241 | ['F', 'I', 'L'] | HTT |
| 172 | 514 | ['F', 'L'] | TTW |
| 174 | 520 | ['L', 'Y'] | TAT, CTG |
| 175 | 523 | ['A', 'G', 'M', 'S'] | ATG, RGT, GCA |
| 177 | 529 | ['A', 'S', 'T'] | DCA |
| 179 | 535 | ['A', 'F', 'L', 'M'] | GCA, MTG, TTT |

*counting from the beginning of *Mth*UPO sequence with signal peptide.

**Table S8** Sequence space of MLEE-enriched library

| Pos. | WT | Mutations |
|---|---|---|
| 59 | F | KTI |
| 63 | F | LI |
| 156 | Y | L |
| 159 | S | A |
| 161 | A | FL |
| **Seq. space** | | **144** |

**Table S9** Ambiguous codons at each mutated position for the MLEE-enriched library.

| Protein Position* | DNA_position* | Encoded amino acids | Ambiguous codons |
|---|---|---|---|
| 77 | 229 | ['F', 'I', 'K', 'T'] | WTT, AMA |
| 81 | 241 | ['F', 'I', 'L'] | HTT |
| 174 | 520 | ['L', 'Y'] | TAT, CTG |
| 177 | 529 | ['A', 'S'] | KCA |
| 179 | 535 | ['A', 'F', 'L'] | GCA, TTW |

*Counting from the beginning of *Mth*UPO sequence with signal peptide.

**Table S10** Oligonucleotides sequences for variable fragments used at initial round

| Pos | Name | Sequence |
|---|---|---|
| 59, 63. | F2_WT | CAACAATTTCGGTTTGTAGGTCTCAACAAAACCTTAGCCAGC**TTT**CTGTTTGAC**TTT**GCATTAACAACGAATCCGATGAGACCAGCGCTCTTACTTTAAA |
| | F2_V1 | CAACAATTTCGGTTTGTAGGTCTCAACAAAACCTTAGCCAGC**WTT**CTGTTTGAC**HTT**GCATTAACAACGAATCCGATGAGACCAGCGCTCTTACTTTAAA |
| | F2_V2 | CAACAATTTCGGTTTGTAGGTCTCAACAAAACCTTAGCCAGC**AMA**CTGTTTGAC**HTT**GCATTAACAACGAATCCGATGAGACCAGCGCTCTTACTTTAAA |
| 154, 156, 157, 159, 161. | F4_WT | *C*AACAATTTCGGTTTGTAGGTCTCAGGAGATGCT**TTC**ACA**TATGGG**GAA**TCT**GCT**GCG**TATGTGGTAGTGTTAGGTTGAGACCAGCGCTCTTACTTTAAA |
| | F4_V1 | CAACAATTTCGGTTTGTAGGTCTCAGGAGATGCT**TTW**ACA**TATATG**GAA**DCA**GCT**GCA**TATGTGGTAGTGTTAGGTTGAGACCAGCGCTCTTACTTTAAA |
| | F4_V2 | CAACAATTTCGGTTTGTAGGTCTCAGGAGATGCT**TTW**ACA**TATATG**GAA**DCA**GCT**MTG**TATGTGGTAGTGTTAGGTTGAGACCAGCGCTCTTACTTTAAA |
| | F4_V3 | CAACAATTTCGGTTTGTAGGTCTCAGGAGATGCT**TTW**ACA**TATATG**GAA**DCA**GCT**TTT**TATGTGGTAGTGTTAGGTTGAGACCAGCGCTCTTACTTTAAA |
| | F4_V4 | CAACAATTTCGGTTTGTAGGTCTCAGGAGATGCT**TTW**ACA**TATRGT**GAA**DCA**GCT**GCA**TATGTGGTAGTGTTAGGTTGAGACCAGCGCTCTTACTTTAAA |
| | F4_V5 | CAACAATTTCGGTTTGTAGGTCTCAGGAGATGCT**TTW**ACA**TATRGT**GAA**DCA**GCT**MTG**TATGTGGTAGTGTTAGGTTGAGACCAGCGCTCTTACTTTAAA |
| | F4_V6 | CAACAATTTCGGTTTGTAGGTCTCAGGAGATGCT**TTW**ACA**TATRGT**GAA**DCA**GCT**TTT**TATGTGGTAGTGTTAGGTTGAGACCAGCGCTCTTACTTTAAA |
| | F4_V7 | CAACAATTTCGGTTTGTAGGTCTCAGGAGATGCT**TTW**ACA**TATGCA**GAA**DCA**GCT**GCA**TATGTGGTAGTGTTAGGTTGAGACCAGCGCTCTTACTTTAAA |
| | F4_V8 | CAACAATTTCGGTTTGTAGGTCTCAGGAGATGCT**TTW**ACA**TATGCA**GAA**DCA**GCT**MTG**TATGTGGTAGTGTTAGGTTGAGACCAGCGCTCTTACTTTAAA |
| | F4_V9 | CAACAATTTCGGTTTGTAGGTCTCAGGAGATGCT**TTW**ACA**TATGCA**GAA**DCA**GCT**TTT**TATGTGGTAGTGTTAGGTTGAGACCAGCGCTCTTACTTTAAA |

**(Continues)**

**(Continues)**

| | |
|---|---|
| F4_V10 | CAACAATTTCGGTTTGTAGGTCTCAGGAGATGCT**TTW**ACA**CTGATG**<br>GAA**DCA**GCT**GCA**TATGTGGTAGTGTTAGGTTGAGACCAGCGCTCTT<br>ACTTTAAA |
| F4_V11 | CAACAATTTCGGTTTGTAGGTCTCAGGAGATGCT**TTW**ACA**CTGATG**<br>GAA**DCA**GCT**MTG**TATGTGGTAGTGTTAGGTTGAGACCAGCGCTCTT<br>ACTTTAAA |
| F4_V12 | CAACAATTTCGGTTTGTAGGTCTCAGGAGATGCT**TTW**ACA**CTGATG**<br>GAA**DCA**GCT**TTT**TATGTGGTAGTGTTAGGTTGAGACCAGCGCTCTT<br>ACTTTAAA |
| F4_V13 | CAACAATTTCGGTTTGTAGGTCTCAGGAGATGCT**TTW**ACA**CTGRGT**<br>GAA**DCA**GCT**GCA**TATGTGGTAGTTTAGGTTGAGACCAGCGCTCTTA<br>CTTTAAA |
| F4_V14 | CAACAATTTCGGTTTGTAGGTCTCAGGAGATGCT**TTW**ACA**CTGRGT**<br>GAA**DCA**GCT**MTG**TATGTGGTAGTGTTAGGTTGAGACCAGCGCTCTT<br>ACTTTAAA |
| F4_V15 | CAACAATTTCGGTTTGTAGGTCTCAGGAGATGCT**TTW**ACA**CTGRGT**<br>GAA**DCA**GCT**TTT**TATGTGGTAGTGTTAGGTTGAGACCAGCGCTCTT<br>ACTTTAAA |
| F4_V16 | CAACAATTTCGGTTTGTAGGTCTCAGGAGATGCT**TTW**ACA**CTGGCA**<br>GAA**DCA**GCT**GCA**TATGTGGTAGTGTTAGGTTGAGACCAGCGCTCTT<br>ACTTTAAA |
| F4_V17 | CAACAATTTCGGTTTGTAGGTCTCAGGAGATGCT**TTW**ACA**CTGGCA**<br>GAA**DCA**GCT**MTG**TATGTGGTAGTGTTAGGTTGAGACCAGCGCTCTT<br>ACTTTAAA |
| F4_V18 | CAACAATTTCGGTTTGTAGGTCTCAGGAGATGCT**TTW**ACA**CTGGCA**<br>GAA**DCA**GCT**TTT**TATGTGGTAGTGTTAGGTTGAGACCAGCGCTCTT<br>ACTTTAAA |

**Bald:** codons at mutated positions. Underscore: Adapter sequences for Golden Gate assembly and PCR.

**Table S11** Oligonucleotides sequences for variable fragments used at MLEE-enriched library

| Pos. | Name | Sequence |
|---|---|---|
| 59, 63 | | The same as Table S10 |
| 156, 159, 161 | F4_V19 | CAACAATTTCGGTTTGTAGGTCTCAGGAGATGCTTTCACA**CTG**GGGGAA**KCA**GCT**GCA**TATGTGGTAGTGTTAGGTTGAGACCAGCGCTCTTACTTTAAA |
| | F4_V20 | CAACAATTTCGGTTTGTAGGTCTCAGGAGATGCTTTCACA**TAT**GGGGAA**KCA**GCT**GCA**TATGTGGTAGTGTTAGGTTGAGACCAGCGCTCTTACTTTAAA |
| | F4_V21 | CAACAATTTCGGTTTGTAGGTCTCAGGAGATGCTTTCACA**CTG**GGGGAA**KCA**GCT**TTW**TATGTGGTAGTGTTAGGTTGAGACCAGCGCTCTTACTTTAAA |
| | F4_V22 | CAACAATTTCGGTTTGTAGGTCTCAGGAGATGCTTTCACA**TAT**GGGGAAKCAGCT**TTW**TATGTGGTAGTGTTAGGTTGAGACCAGCGCTCTTACTTTAAA |

**Bald:** codons at mutated positions. Underscore: Adapter sequences for Golden Gate assembly and PCR.

**Table S12** Primers used for library construction

| Name | Sequence | Note |
|---|---|---|
| F1_fwd | TTGGTCTCAAATGTTTGCTTTTTATTTCTTGACTGCTTG | For constant fragments |
| F1_rev | TTGGTCTCATTGTTGATGTGCAATGCTTCGAAC | |
| F3_fwd | TTGGTCTCACCGAAGAATACCTCGACG | |
| F3_rev | TTGGTCTCACTCCTAATTCCGACATGGAATAC | |
| F5_fwd | TTGGTCTCAAGGTGACAAAGAGTCTCGTAC | |
| F5_rev | TTGGTCTCAAAGCTCAAGTAATACCAGCAGC | |
| F2_F4_fwd | CAACAATTTCGGTTTGTAGGTCTCA | For variable fragments |
| F2_F4_rev | TTTAAAGTAAGAGCGCTGGTCTCA | |
| Seq_fwd | CATCTTATTAAAGTATCATCAAGAAATTGTTA | Sequencing |
| Seq_rev | AAAACGAACTAACTAATGTTTAAGTAAAAGAA | |

**Table S13** GC-MS setup for screening and TON determination

| Experiment | GC-MS | Split | Temperature program |
|---|---|---|---|
| MISER | Shimazu GCMS QP2010 Ultra | 25 | 220 °C 130 min;<br>20 °C/min to 300 °C, hold 30 min |
| TON | Shimazu GCMS QP2020 NX | 5 | 110 °C;<br>3.5 °C/min to 170°C, hold 3 min;<br>50 °C/min to 300°C, hold 3 min |

**Table S14** Primers used for multiplex NGS

| Name* | sequence | barcode |
|---|---|---|
| CLM_1_f | ACACTCTTTCCCTACACGACGCTCTTCCGATCTGATCATGCACCCAAGACCACTCTCCGG | GATCATG |
| CLM_2_f | ACACTCTTTCCCTACACGACGCTCTTCCGATCTTACATGGCACCCAAGACCACTCTCCGG | TACATGG |
| CLM_3_f | ACACTCTTTCCCTACACGACGCTCTTCCGATCTAAGCACCCACCCAAGACCACTCTCCGG | AAGCACC |
| CLM_4_f | ACACTCTTTCCCTACACGACGCTCTTCCGATCTTGGCTCACACCCAAGACCACTCTCCGG | TGGCTCA |
| CLM_5_f | ACACTCTTTCCCTACACGACGCTCTTCCGATCTCTTGCTCCACCCAAGACCACTCTCCGG | CTTGCTC |
| CLM_6_f | ACACTCTTTCCCTACACGACGCTCTTCCGATCTGAAGCGTCACCCAAGACCACTCTCCGG | GAAGCGT |
| CLM_7_f | ACACTCTTTCCCTACACGACGCTCTTCCGATCTTCTCCATCACCCAAGACCACTCTCCGG | TCTCCAT |
| CLM_8_f | ACACTCTTTCCCTACACGACGCTCTTCCGATCTTTGAAGGCACCCAAGACCACTCTCCGG | TTGAAGG |
| CLM_9_f | ACACTCTTTCCCTACACGACGCTCTTCCGATCTGAATGTCCACCCAAGACCACTCTCCGG | GAATGTC |
| CLM_10_f | ACACTCTTTCCCTACACGACGCTCTTCCGATCTATCTCCACACCCAAGACCACTCTCCGG | ATCTCCA |
| CLM_11_f | ACACTCTTTCCCTACACGACGCTCTTCCGATCTGCGTTATCACCCAAGACCACTCTCCGG | GCGTTAT |
| CLM_12_f | ACACTCTTTCCCTACACGACGCTCTTCCGATCTTGCACCACACCCAAGACCACTCTCCGG | TGCACCA |
| CLM_13_f | ACACTCTTTCCCTACACGACGCTCTTCCGATCTTGCCTATCACCCAAGACCACTCTCCGG | TGCCTAT |
| CLM_14_f | ACACTCTTTCCCTACACGACGCTCTTCCGATCTAGGAATCCACCCAAGACCACTCTCCGG | AGGAATC |
| CLM_15_f | ACACTCTTTCCCTACACGACGCTCTTCCGATCTTCCACTGCACCCAAGACCACTCTCCGG | TCCACTG |
| CLM_16_f | ACACTCTTTCCCTACACGACGCTCTTCCGATCTTTGTACCCACCCAAGACCACTCTCCGG | TTGTACC |
| CLM_17_f | ACACTCTTTCCCTACACGACGCTCTTCCGATCTTTCGAGTCACCCAAGACCACTCTCCGG | TTCGAGT |
| CLM_18_f | ACACTCTTTCCCTACACGACGCTCTTCCGATCTCTTCAGCCACCCAAGACCACTCTCCGG | CTTCAGC |
| CLM_19_f | ACACTCTTTCCCTACACGACGCTCTTCCGATCTCAGTGCACACCCAAGACCACTCTCCGG | CAGTGCA |
| CLM_20_f | ACACTCTTTCCCTACACGACGCTCTTCCGATCTTGCTGTCCACCCAAGACCACTCTCCGG | TGCTGTC |
| CLM_21_f | ACACTCTTTCCCTACACGACGCTCTTCCGATCTCGCCATTCACCCAAGACCACTCTCCGG | CGCCATT |
| CLM_22_f | ACACTCTTTCCCTACACGACGCTCTTCCGATCTGCCATGACACCCAAGACCACTCTCCGG | GCCATGA |
| CLM_23_f | ACACTCTTTCCCTACACGACGCTCTTCCGATCTCACAACGCACCCAAGACCACTCTCCGG | CACAACG |
| CLM_24_f | ACACTCTTTCCCTACACGACGCTCTTCCGATCTCTTCGCTCACCCAAGACCACTCTCCGG | CTTCGCT |
| ROW_1_r | GACTGGAGTTCAGACGTGTGCTCTTCCGATCTGAACTGCCGGTGTGCGAAGTAGGTGC | GAACTGC |
| ROW_2_r | GACTGGAGTTCAGACGTGTGCTCTTCCGATCTACCAGGTCGGTGTGCGAAGTAGGTGC | ACCAGGT |

**(continues)**

**(continues)**

| | | |
|---|---|---|
| ROW_3_r | GACTGGAGTTCAGACGTGTGCTCTTCCGATCTTCTAGAGCGGTGTGCGAAGTAGGTGC | TCTAGAG |
| ROW_4_r | GACTGGAGTTCAGACGTGTGCTCTTCCGATCTCACACAACGGTGTGCGAAGTAGGTGC | CACACAA |
| ROW_5_r | GACTGGAGTTCAGACGTGTGCTCTTCCGATCTGTGGAACCGGTGTGCGAAGTAGGTGC | GTGGAAC |
| ROW_6_r | GACTGGAGTTCAGACGTGTGCTCTTCCGATCTATATGCCCGGTGTGCGAAGTAGGTGC | ATATGCC |
| ROW_7_r | GACTGGAGTTCAGACGTGTGCTCTTCCGATCTGGTCTGACGGTGTGCGAAGTAGGTGC | GGTCTGA |
| ROW_8_r | GACTGGAGTTCAGACGTGTGCTCTTCCGATCTGTGAGATCGGTGTGCGAAGTAGGTGC | GTGAGAT |
| ROW_9_r | GACTGGAGTTCAGACGTGTGCTCTTCCGATCTTTGGCAGCGGTGTGCGAAGTAGGTGC | TTGGCAG |
| ROW_10_r | GACTGGAGTTCAGACGTGTGCTCTTCCGATCTATGCCTGCGGTGTGCGAAGTAGGTGC | ATGCCTG |
| ROW_11_r | GACTGGAGTTCAGACGTGTGCTCTTCCGATCTTCCGAAGCGGTGTGCGAAGTAGGTGC | TCCGAAG |
| ROW_12_r | GACTGGAGTTCAGACGTGTGCTCTTCCGATCTGGCTTACCGGTGTGCGAAGTAGGTGC | GGCTTAC |
| ROW_13_r | GACTGGAGTTCAGACGTGTGCTCTTCCGATCTAGTTGGCCGGTGTGCGAAGTAGGTGC | AGTTGGC |
| ROW_14_r | GACTGGAGTTCAGACGTGTGCTCTTCCGATCTAACGATGCGGTGTGCGAAGTAGGTGC | AACGATG |
| ROW_15_r | GACTGGAGTTCAGACGTGTGCTCTTCCGATCTACTACCGCGGTGTGCGAAGTAGGTGC | ACTACCG |
| ROW_16_r | GACTGGAGTTCAGACGTGTGCTCTTCCGATCTGGTGTCTCGGTGTGCGAAGTAGGTGC | GGTGTCT |
| NGS_inner_fwd | CACCCAAGACCACTCTCCGGACGCCTTGTTCGAAGCATTG | |
| NGS_inner_rev | CGGTGTGCGAAGTAGGTGCTGACAGTACGAGACTCTTTGTC | |

***barcode_1**: CLM_1_F to CLM_12_F as column indicator, Rrow_1_R to ROW_8_R as row indicator;

**barcode_2**: CLM_13_F to CLM_24_F as column indicator, Rrow_1_R and ROW_8_R as row indicator;

**barcode_3**: CLM_1_F to CLM_12_F as column indicator, Rrow_9_R to ROW_16_R as row indicator;

**barcode_4:** CLM_13_F to CLM_24_F as column indicator, Rrow_9_R to ROW_16_R as row indicator

**Table S15 Hyperparameters optimization for supervised ML**

| Hyperparameters | Grid |
|---|---|
| mlp_hidden_layer_sizes | (22), (22, 11), (22,11, 5)<br>(77), (77, 38), (77, 38, 19) |
| mlp_activation | tanh, relu |
| mlp_alpha | 1e-4, 1e-3, 1e-2 |
| mlp_learning_rate_init | 0.001, 0.005, 0.01 |
| mlp_weight_exp | 0, 0.5, 1, 1.5 |
